# Niche-driven phenotypic plasticity and *cis*-regulatory dynamics of a revised model for intestinal secretory differentiation

**DOI:** 10.1101/2025.02.07.636917

**Authors:** Swarnabh Bhattacharya, Guodong Tie, Pratik N. P. Singh, Ermanno Malagola, Onur Eskiocak, Ruiyang He, Judith Kraiczy, Wei Gu, Yakov Perlov, Semir Beyaz, Timothy C. Wang, Qiao Zhou, Ramesh A. Shivdasani

## Abstract

**HIGHLIGHTS:**

- Delineation of chromatin and mRNA dynamics of intestinal secretory differentiation
- Paneth cells have few unique enhancers and share mRNAs and TFs with goblet cells
- Unlike other secretory derivatives, goblet and Paneth cells are not specified *per se*
- Niche factors, especially BMP signaling, define goblet and Paneth phenotypes

Enterocytes and four secretory cell types derive from stem cells located in intestinal crypts. Whereas secretory goblet and Paneth cells have long been considered distinct, we find high overlap in their transcripts and sites of accessible chromatin, in marked contrast to those of sibling enteroendocrine or tuft cells. Mouse and human goblet and Paneth cells express extraordinary fractions of selective antimicrobial genes, reflecting specific and variable gene responses to local niche signals. Wnt signaling retains few ATOH1^+^ secretory daughters in crypt bottoms, where an absence of BMP signaling potently induces Paneth features; those that move away from crypt bottoms acquire classic goblet properties. These post-mitotic cellular phenotypes and their underlying accessible *cis*-elements interconvert readily. Thus, goblet and Paneth properties represent alternative manifestations of a single versatile signal-responsive secretory cell. These findings reveal exquisite niche-dependent cell plasticity and the *cis*-regulatory dynamics of an updated unitarian model of the intestinal epithelial lineage.

## INTRODUCTION

For half a century, understanding of the intestinal epithelium has rested on a ‘unitarian’ model in which multipotent stem cells continuously generate absorptive enterocytes and distinct secretory (goblet, enteroendocrine (EE), Paneth) cells.^1,2^ Lgr5^+^ cells in crypt bottoms were later identified as intestinal stem cells (ISCs)^3^ and tuft cells as an additional secretory type (Figure 1A).^4,5^ Importantly, ISC attributes are not cell-autonomous, but are conferred on epithelial cells in the crypt base by highly localized mesenchymal cues, especially Wnt/Rspondin (Rspo) agonists and BMP antagonists.^6–9^ As ISC progeny moves away from the base, restricted transcription factors (TFs) drive their acquisition of distinctive gene expression profiles and cell characteristics. First, ATOH1 demarcates secretory from enterocyte progenitors.^10,11^ This pivotal divergence belies the observation that goblet cells and enterocytes share a substantial fraction of *cis*-regulatory elements (mainly enhancers) with active histone marks and open chromatin.^12^ Subsequently, daughter cells with high expression of NEUROG3 or POU2F3 adopt EE or tuft cell fates, respectively.^13–16^ NEUROG3^+^ progenitors, for example, activate *cis*-elements that are selectively accessible in EE cell chromatin.^17,18^ Antimicrobial Paneth cells, which are uniquely long-lived and reside among ISCs in the crypt base, are ostensibly as specialized as EE or tuft cells and various perturbations are reported to affect their numbers or morphology. For example, Paneth cells are deficient or absent in intestines that lack TF genes *Gfi1* or *Sox9* or carry histone lysine-to-methionine mutations; however, no implicated factor is exclusive to –or necessarily expressed in– Paneth cells and the mutant mice generally also show goblet cell defects.^19–24^

**Figure 1.**
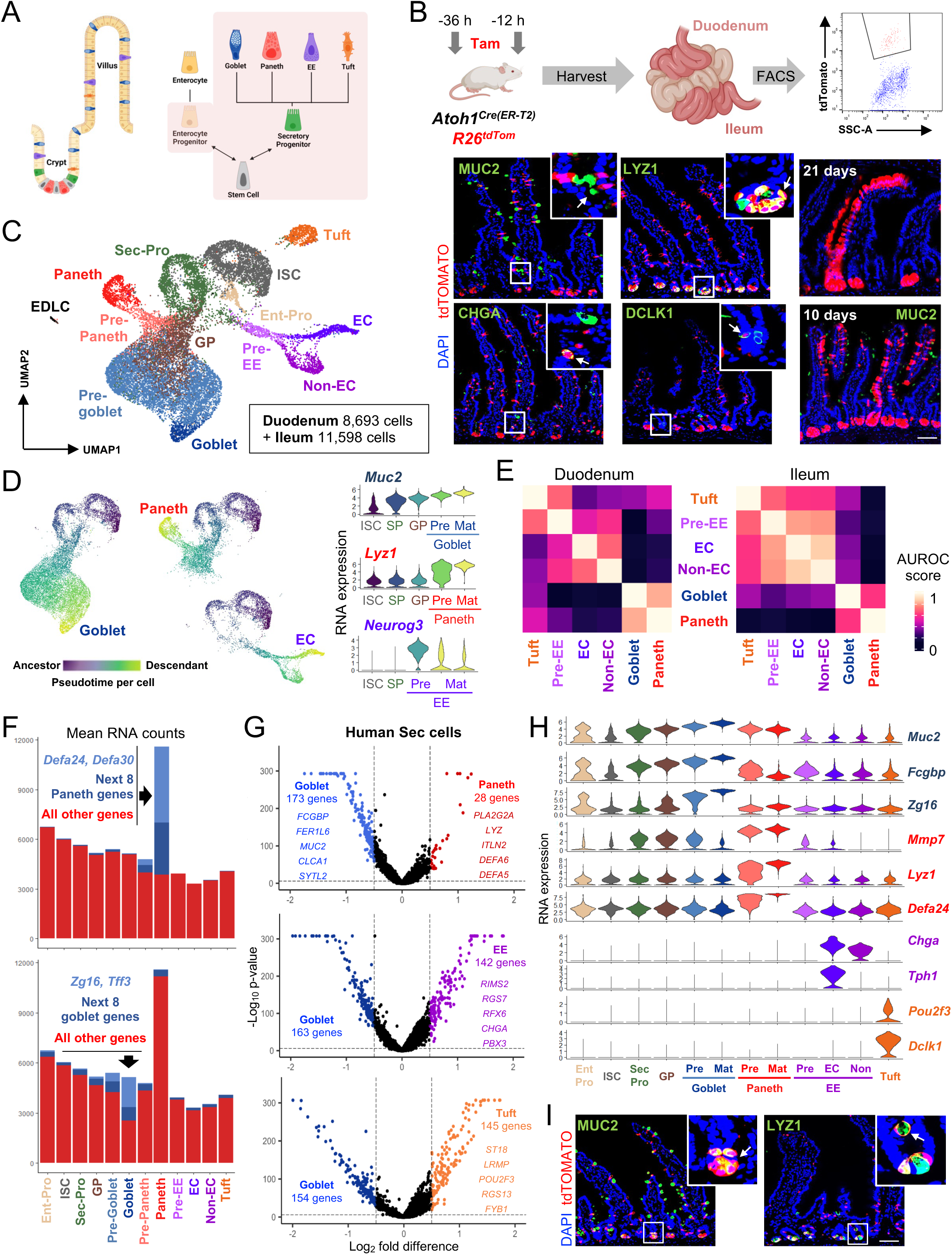
High transcriptional overlap between goblet and Paneth cells. **(A)** Unitarian model of intestinal epithelial self-renewal. Boxes demarcate the cells expected to be labelled in *Atoh1^Cre-ER(T2)^;R26^Tom^* mice. **(B)** Lineage tracing and cell isolation from *Atoh1^Cre-ER(T2)^;R26^Tom^* mice treated with 2 doses of tamoxifen (Tam). Duodenal and ileal crypt cells were harvested 12 h after the second dose and a representative duodenal tdTomato FACS plot is shown. Micrographs reveal efficient labeling of all secretory cell types, abundant ISC progeny at 10 days, and occasional ISC-derived clonal ‘ribbons’ at 21 days. n=3 mice, scale bar 50 µm. **(C)** Integrated UMAP plot from scRNA analysis of 20,291 combined duodenal (n=2 mice) and ileal (n=2 mice) Tom^+^ cells, showing each expected population. **(D)** Slingshot pseudotime trajectories for mature goblet, Paneth, and enterochromaffin (EC) cells, showing their common origins in ISCs and secretory progenitors (Sec-Pro). Only cells that contribute to a trajectory acquire scores (colour) and dark-to-light color gradients depict pseudotime progression. Cell-enriched marker transcripts support precursor (Pre) and mature (Mat) cell assignments. **(E)** MetaNeighbour area under the receiver operating characteristic (AUROC) analysis, showing high similarity of goblet and Paneth, compared to EE and tuft, cells. **(F)** Mean RNA counts in duodenal cell types, showing extreme skewing in Paneth cells and disproportionate enrichment of a few select genes in Paneth and goblet cells. **(G)** Genes differentially expressed (log_2_ fold-difference >0.5, FDR <0.05 determined by the Wald test) in human goblet and other secretory cell types.^33^ Genes with the largest differences are named. **(H)** Pseudobulk-averaged expression of conventional secretory cell markers. Highly overlapping expression in goblet and Paneth cells contrasts with selective expression of EE and tuft markers. **(I)** Representative micrographs of MUC2 and LYZ1 immunostaining in *Atoh1^Cre-ER(T2)^;R26^Tom^* intestines. Although MUC2 and LYZ1 are highest in goblet and Paneth cells, respectively, weaker signals are apparent in Paneth and rare goblet cells, respectively. n=5 mice, scale bar 50 µm. See also Figures S1 and S2.

To identify TFs that might specify Paneth cells, we examined mRNA and chromatin in the derivatives of *Atoh1^+^* secretory progenitors. No TF genes and few *cis*-elements distinguished Paneth from goblet cells. Rather, in mice and in primary mouse and human ISC-derived cultures, goblet- and Paneth-enriched genes were highly and reciprocally responsive to Wnt and especially BMP signaling. EE and tuft cells cease to express *Atoh1* after they are specified by other downstream TFs, and their chromatin signatures resist significant modulation by Wnt or BMP signaling. In contrast, *Atoh1* remains expressed in the dominant secretory derivative, where it is required to maintain goblet or Paneth properties. Thus, localized niche signals act on a shared permissive epigenome to drive alternative goblet or Paneth features in a single ATOH1-dependent secretory cell type, and Paneth properties reflect the high-Wnt and low-BMP milieu they co-inhabit with crypt base ISCs. Together, these findings refine the unitarian model, define its *cis*-regulatory basis, demonstrate exquisite secretory cell sensitivity –on a par with that of ISCs– to sub-epithelial signals, and further implicate the mesenchymal niche in instructing epithelial cell character.

## RESULTS

### Unusual and overlapping features of goblet and Paneth cell transcriptomes

To capture emerging intestinal secretory cells, we injected adult *Atoh1^Cre(ER-T^*^2^*^)^*;*Rosa26^L-S-L-tdTomato^* (*Atoh1^Cre^*;*R26^Tom^*) mice^25,26^ with two doses of tamoxifen 24 h apart and isolated tdTom^+^ crypt cells by flow cytometry 12 h after the second dose (Figures 1B and S1A). At this time, tdTom stained nearly every villus MUC2^+^ cell with typical goblet morphology, LYZ1^+^ Paneth cells in the crypt base, and small fractions of CHGA^+^ EE and DCLK1^+^ tuft cells (Figure 1B), indicating that terminal goblet and Paneth cells express *Atoh1*, but mature EE and tuft cells do not. Rather, tdTom^+^ EE and tuft cells and MUC2^+^ goblet cells recently derived from *Atoh1^+^* secretory progenitors were confined to crypts (Figure 1B). These progenitors are known to revert at some frequency into multipotential ISCs,^27^ but few such ISCs become clonally fixed.^28,29^ Accordingly, 10 days after tamoxifen exposure, before clonal fixation, numerous tdTom^+^ villus enterocytes reflected the activity of transient *Atoh1^+^* cell-derived ISCs, but by 21 days only 2 to ∼5 crypts and adjoining villi were fully tdTom^+^ in duodenal and ileal cross-sections (Figures 1B and S1B). From crypts we therefore expected to recover tdTom^+^ ISCs, secretory progenitors, precursors of each secretory type, Paneth cells, some mature goblet cells, and as a consequence of ISC labeling, some tdTom^+^ enterocyte progenitors (Figure 1A, shaded box). mRNA profiles of >20,000 single tdTom^+^ cells isolated from duodenal or ileal crypts 12 h and 36 h after Cre activation, coupled with marker-based cell assignments, identified each of these populations (Figures S1C-E). Cellular trajectories imputed by Slingshot^30^ or RNA velocity^31^ traced the origin of differentiated cells in each region from ISCs via secretory progenitors and specific precursors (Figure S1F). For ease of presentation, we merged duodenal and ileal cells (Figure 1C), which showed similar population and ontogenic structures. Canonical secretory markers validated the assignments of goblet, Paneth, and EE precursor (pre) cells, which expressed classic markers at levels lower than their terminal descendants (Figure 1D). Tuft and EE cells clustered apart from the others, as did epithelium-derived lymphoid-like cells (EDLCs – characterized in a companion manuscript), and pre-EE cells branched as expected into serotonergic enterochromaffin and peptide hormone-producing (non-enterochromaffin) subtypes.

Goblet and Paneth cell transcriptomes resembled each other more than any other pair of cell types, roughly to the tune of the two EE cell branches (Figures 1E and S1D), and appeared to descend from a common goblet/Paneth precursor (GP, Figures 1C and 1D). While >500 genes met stringent criteria for differential expression in tuft or EE cells relative to goblet cells, an order of magnitude fewer genes –dominated by *Lyz1*, *Mmp7*, *Itln1*, *Mptx2*, and α-Defensins– were enriched in Paneth cells (Figure S1G, Table S1). Moreover, mean scRNA counts ranged from 2,748 and 6,738 in other cell types, but Paneth cells averaged 11,587 (duodenum) to 16,292 (ileum) RNA reads per cell (Figures 1F and S2A). Ten transcripts together accounted for 66.6% (duodenum) to 72.4% (ileum) of all Paneth cell mRNA and just one (ileum) or two (duodenum) transcripts, *Defa24* and *Defa30*, gave more reads than all transcripts combined in other cells (Figures 1F and S2B). Goblet cell RNA counts were in the same range as other secretory cells, but the top 10 genes –including *Zg16*, *Muc2*, *Tff3* and *Clca1*– represented 50.4% of the transcriptome; goblet and Paneth precursors lacked this extreme transcriptional skewing and the 10 dominant transcripts in tuft or EE cells represented 9.9 to 13.5% of all mRNAs (Figures 1F, S2A, and S2B). Mouse goblet and Paneth cells captured in a previous scRNA study^32^ affirm the disproportionate contributions of a few genes to atypically high mRNA loads (Figure S2C). Human Paneth cells^33^ also show limited distinction from goblet cells (Figures 1G and S2D and Table S1).

The sizable fraction of RNAs encoding a few genes affects normalization of RNA counts and exaggerates differences across cell types; when comparing genes, we therefore excluded those with >200 RNA reads per cell, except the gene under interrogation (Figure S2E). Whereas tuft and EE cell markers were tightly restricted, ISCs, enterocytes, and all secretory derivatives expressed appreciable levels of classic goblet and Paneth markers (Figure 1H). Terminal goblet and Paneth cells had about the same levels of each other’s markers as did their presumptive GP precursors (e.g., *Muc2* and *Zg16* in Paneth or *Lyz1* and *Mmp7* in goblet), and cell-enriched markers increased in their respective precursors. The most notable distinction was massive Paneth cell up-regulation of Defensins, *Lyz1*, *Itln1*, *Mmp7*, and other select genes that goblet cells express at lower levels (Figure 1H); for example, *Defa24* is 150 times higher in Paneth cells, but also the 9^th^ most abundant transcript in ileal crypt goblet cells (Figure S2B). Ambient RNA artifacts cannot explain the sum of these findings and, although MUC2 and LYZ1 proteins dominate in goblet and Paneth cells, respectively, each is present in the other (Figure 1I). Moreover, a Paneth-specific gene module expresses at higher levels per cell in all ileal than in duodenal cell types (Figure S2F). In mice infected with *Salmonella typhimurium* or with the helminth *Strongyloides venezuelensis* and in mice treated with infection mimetics IL-25 or succinate, LYZ1 and MMP7 proteins were no longer confined to Paneth cells at the base of distal intestinal crypt but extended well into crypts, in mature cells with goblet morphology (Figure 2A). Indeed, scRNA-seq data^32^ from mice infected with *Salmonella* or with the helminth *H. polygyrus* confirmed broad expression of Paneth-selective mRNAs in many epithelial –especially goblet– cells (Figure S2G). Thus, much of the Paneth cell transcriptome represents a specific microbe-responsive program and goblet and Paneth cells may represent phenotypic variants of one secretory type that differ principally in expressing specific antimicrobial genes.

**Figure 2.**
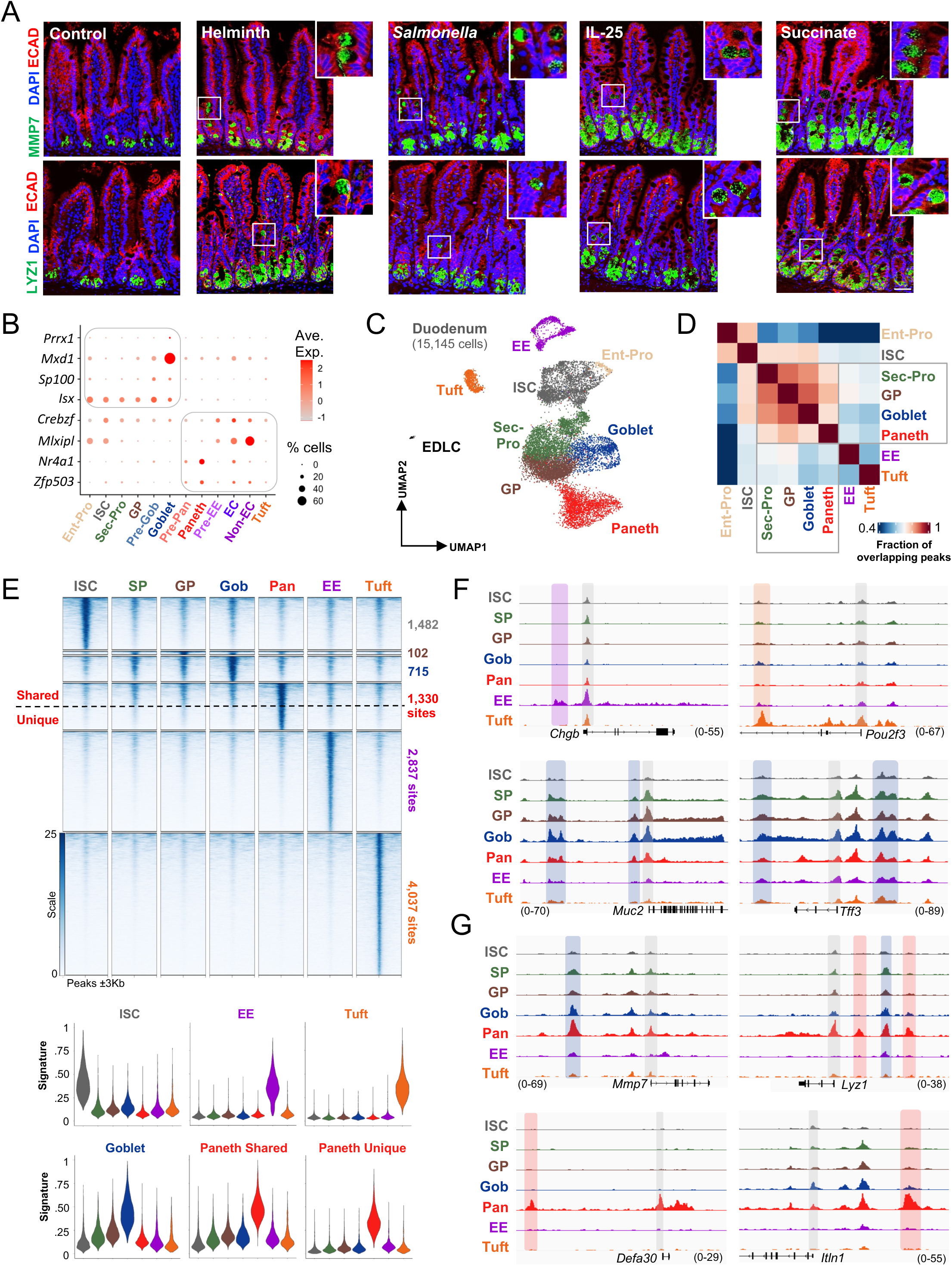
Shared *cis*-element profile and lack of unique TFs in goblet and Paneth cells. **(A)** Representative micrographs of ECAD-, MMP7-, and LYZ1-immunostained ileal tissue from uninfected control, *S. venezuelensis* (helminth – 14 days) infected, *Salmonella typhimurium* infected (5 days), IL-25 treated (7 days) or succinate treated (7 days) mice. Paneth markers MMP7 and LYZ1 in each case extend well into crypts and in mature cells with goblet morphology (insets). n=3 mice, scale bar 50 µm. **(B)** Few TF transcripts are enriched in goblet (villus Tom^+^ cells 36 h after tamoxifen injection) vs. Paneth (crypt Tom^+^ cells 12 days after tamoxifen injection) cells. Even those few TFs are expressed in other secretory cell types and only in small fractions of single cells. **(C)** UMAP plot from snATAC analysis of 15,145 duodenal Tom^+^ crypt cells (n=2 mice injected with tamoxifen 24 h apart), showing *cis*-element neighborhoods for secretory cell populations. **(D)** Fractions of overlapping MACS2-called peaks in individual pseudobulk populations. Global open chromatin was similar in secretory progenitors, goblet-Paneth precursors (GP), and goblet cells, followed closely by Paneth cells, whereas EE and tuft cell profiles diverged. **(E)** Sites of differential open chromatin enriched in any cell cluster (pseudobulk) over all others (pct.min >0.01, log_2_ fold-difference >0.25, n: 10,505 sites). Paneth-enriched regions are further divided into shared (broadly accessible, n: 665) and unique (uniquely accessible, n: 665). Aggregate ATAC module scores for differentially enriched sites are shown below. **(F-G)** Representative IGV tracks from pseudobulk snATAC at classic EE, tuft, goblet and Paneth cell loci illustrate shared accessibility at goblet and shared Paneth sites and restricted accessibility at EE, tuft, and unique Paneth sites. Promoters are shaded in grey and distance enhancers are shaded in colors corresponding to each secretory cell type. See also Figures S3 and S4.

### Limited transcriptional and chromatin diversity between goblet and Paneth cells

Distinct cell types express unique TF combinations. To identify TFs that might underlie secretory diversity, we examined differential TF mRNA expression after normalizing for overrepresented genes (>200 RNA counts per cell – Figure S2E). Certain TFs were enriched in ISCs/Enterocytes (e.g., *Nr3c1*, *Nfib*, *Hnf4g*) or exclusive to tuft (e.g., *Spib*, *Pou2f3*, *Runx1*) or EE (e.g., *Fev*, *Arx*, *Nkx2-2*, *Neurog3*) cells, as expected,^34–36^ but none were restricted to goblet or Paneth cells (Figure S3A). Because scRNA-seq might miss low-expressed genes, we performed bulk RNA-seq on terminal tdTom^+^ Paneth (from crypts 12 days after tamoxifen) and goblet cells from *Atoh1^Cre^*;*R26^Tom^* mouse villi 1.5 days post-tamoxifen. The few TFs modestly enriched in one or the other (Figure S3B) were also expressed in other cells (Figure 2B) and none matched the specificity of *Neurog3* or *Pou2f3*. Thus, no TF seems to account alone for goblet/Paneth cell divergence. Indeed, whereas *in vivo* loss of the latter TFs eliminates EE or tuft cells, respectively,^13,37^ loss of TFs such as *Gfi1*, *Spdef, Sox9,* or *Bhlha15* affects terminal features or relative proportions of goblet and Paneth cells, but not their presence *per se*.^19–24,38,39^ *Atoh1* is one of few TF transcripts that persist in goblet and Paneth cells after secretory specification (Figure S3C, see Figure 1B). *Atoh1*-independent tuft cells have been reported.^40–42^ To test *Atoh1* requirement in other secretory cells, first, we crossed a floxed null allele^43^ onto the EE-selective *Neurog3^Cre(ER-T^*^2^*^)^* background.^44^ CHGA^+^ EE cell numbers were unaffected after tamoxifen-induced EE-restricted *Atoh1* loss (Figure S3D). Next, we found that after tamoxifen exposure, tdTom^+^ (*Atoh1*^-/-^) cells in *Atoh1^Cre/Fl^*;*R26^Tom^* mice retained CHGA^+^ EE and DLCK1^+^ tuft cells but lacked MUC2 or goblet morphology, as expected from goblet cell conversion to enterocytes;^10^ Paneth morphology and LYZ1 were also absent in crypt bottoms for up to 10 days, when new (tdTom^-^) Paneth cells had appeared (Figure S3E). Thus, consistent with their transcriptional overlap and unlike tuft or EE cells, both terminal goblet and Paneth features require *Atoh1*.

Beyond expressing specific TFs, distinct cell types activate many unique enhancers.^45,46^ To identify potentially unique goblet or Paneth *cis*-elements, we profiled accessible chromatin^47^ at single-nucleus (sn) resolution^48^ in >15,000 tdTom^+^ duodenal crypt cells isolated 12 h and 36 h after Cre induction. Although promoters generally vary little across cells,^46^ hundreds of EE, tuft, and Paneth cell-enriched promoters (>-1 kb and >2 kb from transcription start sites, TSSs) were only accessible in specific graph-based snATAC-seq clusters (Figure S3F). Matching expressed RNAs (Figure S1E) with these accessible promoters (Figure S3G) identified distinct populations as ISCs, enterocyte progenitors, secretory progenitors, EE, tuft, and EDL cells (Figure 2C). Affirming the ISC designation, sites accessible in that cell cluster (pseudobulk) matched those from bulk ATAC-seq of purified Lgr5^+^ ISCs (Figure S3H) and Slingshot^30^ imputed chromatin trajectories from ISCs into discrete cell types via secretory progenitors (Figure S4A). Global profiles of open chromatin were similar in these progenitors, goblet/Paneth precursors, and goblet cells, closely followed by Paneth cells (Figure 2D). About 77,000 sites were comparably accessible in all clusters (Figure S4B); only 8.3% of sites were significantly enriched (log_2_ fold-difference ≥0.25; 1,119 promoters and 9,386 enhancers) in any secretory cell type over all others, yielding *cis*-regulatory signatures for each (Figure 2E and Table S2). EE and tuft signatures were highly restricted, while those for other cell types were only relatively enriched and not exclusive (Figures 2F and S4C).

Paneth-enriched sites fell into two equal groups: shared (accessible elsewhere) or unique to Paneth cells (Figure 2F). Genes abundantly expressed in Paneth cells typically carried multiple open sites, some broadly accessible and others Paneth-restricted; only select loci such as Defensins and *Itln1* had exclusively unique Paneth sites (Figures 2G and S4D). Thus, in contrast to thousands of sites selectively accessible in EE or tuft cells, *cis*-elements are marginally enriched in goblet cells and only a few hundred unique sites distribute across Paneth cell loci that harbor other broadly accessible sites. We confirmed these observations in *Neurog3*-labeled crypt cells, which mainly generate EE cells but, reflecting crypt plasticity, also mark goblet/Paneth precursors.^17,49^ snATAC profiles of unperturbed crypts from a prior study of crypt cell dedifferentiation in *Neurog3-Cre^ER-T^*^2^*;R26^Tom^* mice^17^ confirmed extensive overlap of goblet and Paneth cis-elements and the presence of unique Paneth sites amid broadly accessible ones (Figure S4E). snATAC profiles of human intestinal epithelium^33^ also show unique and shared Paneth *cis*-elements, akin to mouse cells (Figure S4F). Thus, goblet and Paneth epigenomes harbor relatively few differences, further negating their distinction as fixed, TF-specified fates.

### Acute Wnt dependency of the Paneth phenotype without cell re-specification

Consistent with TF-mediated cell specification, EE- or tuft-restricted enhancers were highly enriched for DNA sequence motifs for EE- and tuft-exclusive TFs (Figure 3A and see Figure S3A). In contrast, TF motifs –e.g., ATOH1, AP-1 (JUN/FOS), and FOXA– were modestly enriched at Paneth and goblet *cis*-elements, and only unique Paneth sites were weakly enriched for Wnt-responsive TCF/LEF motifs (Figure 3A). Paneth cells reside in the Wnt-rich crypt base, require Wnt receptor FZD5,^50^ increase in number upon forced activation of canonical Wnt signaling,^51–53^ and mislocalize in *Ephb3^-/-^* and other mouse models.^54–57^ Accordingly, the conventional view is that Wnt signaling “specifies” Paneth cells as a distinct secretory type that homes to crypt bottoms in response to signals such as EphB2/EphB3.^51–53^ We suggest instead that goblet and Paneth cells are different manifestations of the same *Atoh1^+^* secretory cell, which reacts to Wnts by expressing Paneth-enriched genes and to signals far from crypt bottoms by adopting goblet features (Figure 3B).

**Figure 3.**
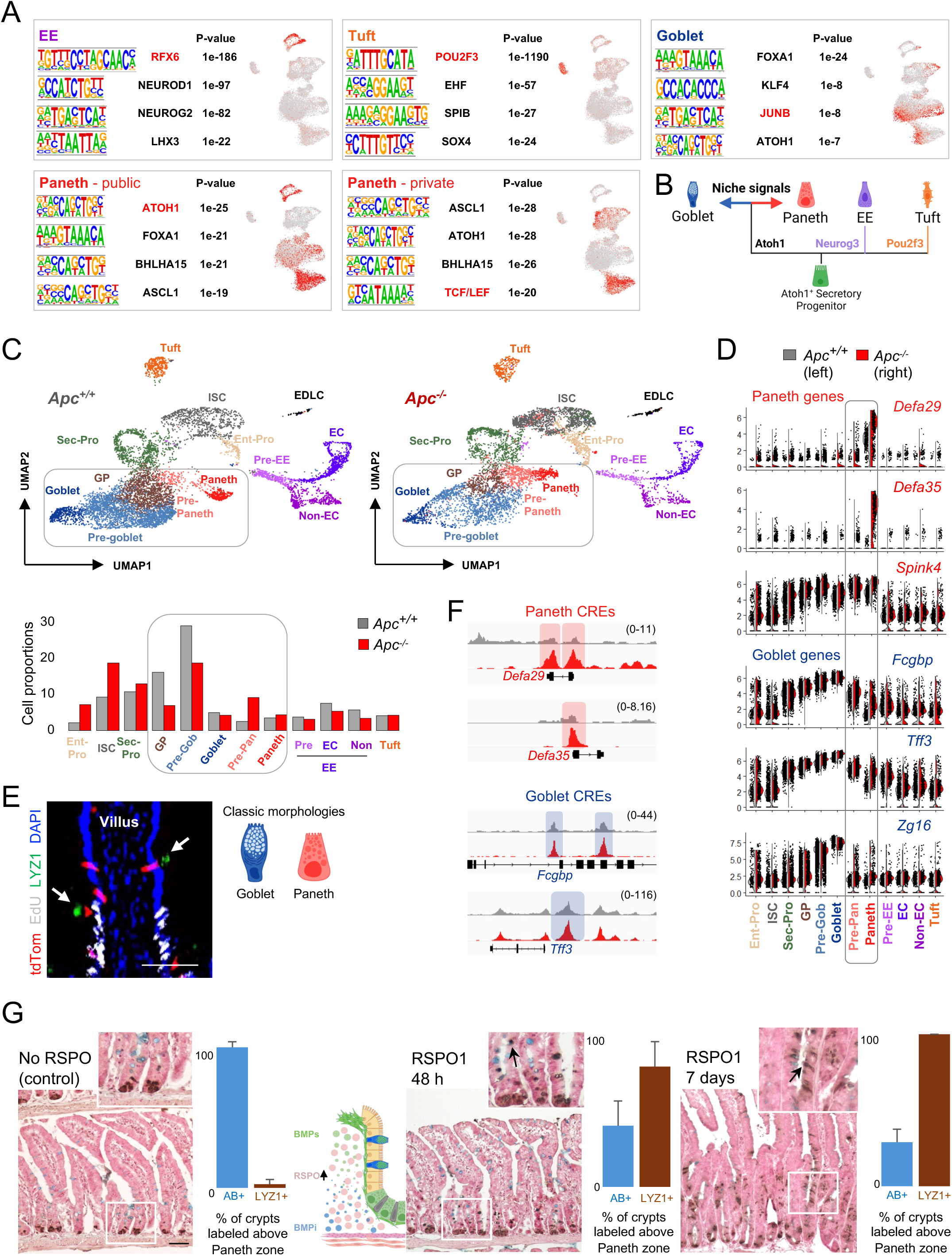
Wnt signaling triggers a phenotypic switch to Paneth features. **(A)** TF-binding sequence motifs enriched at sites differentially accessible in secretory cell types. Enrichment of motifs marked in red is projected on the snATAC UMAP (Figure 2C). **(B)** Hypothesis: goblet and Paneth are minimally different, labile manifestations of Atoh1-determined secretory cells, distinguished by specific niche signals, in contrast to TF-driven specification of EE and tuft cells. **(C)** Merged UMAP plots from scRNA analysis of 15,416 duodenal Tom^+^ cells from wild-type (*Apc^+/+^*, n=2 mice, 8693 cells) and *Apc^-/-^* (n=2 mice, 6723 cells) intestines. Each cell population is indicated and quantified in the graph below. Mice received 2 doses of tamoxifen 24 h apart and tissues were harvested 12 h after the second dose. ISCs and enterocyte progenitors (Ent-Pro) were increased in *Apc^-/-^* crypts as expected, pre-Paneth cells increased at the expense of goblet-Paneth precursors (GP) and pre-goblet cells, and other cell types were minimally perturbed. **(D)** Pseudobulk-averaged expression of classic Paneth (*Defa29*, *Defa35*, *Spink4*) and goblet (*Zg16*, *Fcgbp*, *Tff3*) transcripts, showing elevated α-Defensin expression in all *Apc^-/-^* (including Paneth) cells. Residual goblet-cell transcripts in *Apc^-/-^* Paneth cells indicate conversion from goblet to Paneth features. **(E)** Forced Wnt activation (*Apc* deletion) in *Atoh1^Cre-ER(T2)^;R26^Tom^* mice. Simultaneous single doses of TAM and EdU distinguish pre-existing (above EdU column) from newly generated cells (within EdU^+^ pool). LYZ1 and EdU immunostaining in intestines harvested 24 h later showed increased numbers of ectopic LYZ1^+^ cells, many with goblet morphology (arrows – see adjoining diagrams). n=5 mice of each genotype, scale bar 50 µm. **(F)** Representative IGV tracks (snATAC) from pseudobulk cell populations at goblet and Paneth marker loci in *Apc^+/+^* and *Apc^-/-^* Paneth cells, affirming increased chromatin accessibility at Paneth (Wnt-induced) and goblet (residual) loci. **(G)** Adenoviral RSPO expression enlarged crypts, as expected. Within 48 h LYZ1^+^ cell numbers were increased, including in Alcian blue-stained goblet cells present above the crypt base, and by 1 week nearly every supra-basal secretory cell expressed LYZ1. n=3 mice from each treatment and time point, scale bars 50 µm. See also Figure S5.

Paneth cell excess was previously reported upon Wnt activation in all epithelial cells, so the cause could not be identified.^51–53^ To test whether goblet/Paneth phenotypes are labile in response to canonical Wnt signals, we injected *Atoh1^Cre(ER-T^*^2^*^)^*;*Apc^fl/fl^*;*R26^Tom^* mice with tamoxifen on two consecutive days and isolated tdTom^+^ cells 12 h after the second dose. Relative to *Apc^+/+^* controls (Figure S1E, cells merged and re-normalized with *Apc^-/-^* cells to generate *K* nearest-neighbor (KNN) cell clusters), *Apc^-/-^* ISCs were increased as expected. *Apc^-/-^* secretory progenitor, tuft, EE, and mature goblet or Paneth cell numbers were barely affected, while pre-Paneth cells increased 3.6-fold at the expense of GP and pre-goblet cells (Figure 3C). Paneth-cell mRNAs –especially *α-Defensins*– increased in all *Apc^-/-^* cells, especially in those classified as Paneth, and the levels of goblet cell markers in *Apc^-/-^* Paneth cells pointed to their origin in cells with a recent goblet phenotype (Figure 3D). Indeed, LYZ1 was expressed in cells with goblet morphology both within and above the EdU^+^ column of cells that had replicated during tamoxifen treatment (Figure 3E), i.e., in some post-mitotic goblet cells. Moreover, chromatin accessibility determined by snATAC-seq revealed higher relative abundance of cells with a Paneth profile (Figure S5A) and signals at both Paneth and goblet loci were higher in *Apc^-/-^* than in *Apc^+/+^* Paneth cells (Figure 3F).

To activate canonical Wnt signaling by a different method, we injected mice with adenovirus expressing RSPO1, a Wnt co-factor that expands the ISC pool and delays crypt cell migration into villi.^58^ Within 48 h of RSPO1 exposure, ectopic LYZ1^+^ cells were located above the crypt base and their numbers increased further over 7 days (Figure 3G); these LYZ1^+^ cells displayed goblet morphology and stained with Alcian blue (AB), again signifying the appearance of a Paneth feature in existing goblet cells. Thus, forced Wnt signaling skews goblet/Paneth phenotypic duality in accordance with our predictions. “Ectopic” Paneth cells in other settings^52–54,56,59^ might similarly reflect the expression of Paneth-enriched genes in versatile signal-responsive secretory cells.

### Profound and variable Paneth gene sensitivity and cis-element responses to BMP signaling

Pan-epithelial Wnt activation increases Paneth cell numbers mainly in crypts.^51–53^ Indeed, LYZ1 expression in RSPO1-treated mice was largely confined to their expanded crypts (villus goblet cells were spared, Figure 3G). Although we detected LYZ1 in some post-mitotic *Atoh1^Cre^*; *Apc^fl/fl^* goblet cells (Figure 3E), phenotypic switching occurred principally within the EdU^+^ column, i.e., among cells that were recently in crypts (Figure S5B). Moreover, 7 days after tamoxifen injection, elongated crypts in *Vil1-Cre^ER-T^*^2^*;Apc^fl/fl^* intestines were replete with LYZ1^+^ cells, many with goblet morphology, but villus LYZ1 expression was infrequent and limited (Figure S5C). These findings suggest that signals other than Wnt might restrict the Paneth phenotypic domain.

In addition to RSPOs, cells in the sub- and peri-cryptal mesenchyme also secrete BMP antagonists, such as GREM1, to create an ISC-permissive Wnt^hi^ BMP^lo^ milieu^60,61^ (Figure 4A). The BMP effector genes *Id1* and *Id3* are enriched in mouse and human goblet cells, and mouse crypt goblet cells express more BMP receptor transcripts than Paneth cells (Figures 4B and S5D). To delineate the relative contribution of each pathway in driving secretory phenotypes, first we used intestinal organoids.^62^ We treated duodenal crypt-derived organoids with the Notch inhibitor N-(N-(3,5-difluorophenacetyl)-l-alanyl)-S-phenylglycine-butyl ester (DAPT), which expands secretory cells at the expense of enterocytes^63^ and, importantly, curtails new cell production;^12^ we could therefore attribute phenotypic changes to pre-existing cells. Three days of treatment with DAPT alone activated *Atoh1*, substantially induced secretory transcripts, and abrogated cell replication (Figures S5E, S5F, and S5G), as expected. We then exposed organoids for 4 additional days to a cocktail of recombinant (r) BMP2/4/7 or the synthetic Wnt agonist CHIR 99021 (Ref. ^64^); to avoid BMP blockade, we withdrew rNOG, a routine additive in organoid cultures,^62^ during those 4 days. Wnt activation elevated Paneth transcripts 3-to ∼10-fold and reduced goblet markers modestly (Figure 4C, y-axis on log_10_ scale, *Fcgbp* levels declined ∼64-fold). Conversely, rBMPs increased goblet cell genes 4-to ∼10-fold (only *Zg16* rose >100-fold), while suppressing Paneth markers 25-to >500-fold. Immunostaining verified Paneth dominance upon Wnt activation and its abrogation by BMP signaling (Figures 4C and S5H). Thus, BMPs inhibit Paneth genes more potently than Wnt signaling induces them, helping explain why that phenotype is confined to crypt bottoms. Goblet and Paneth markers displayed a wide range of thresholds and responses to different BMP concentrations (Figure 4D), indicating that niche signals do not *determine* alternative cell states in the conventional sense; rather, each state represents a composite of the genes expressed at different signal concentrations.

**Figure 4.**
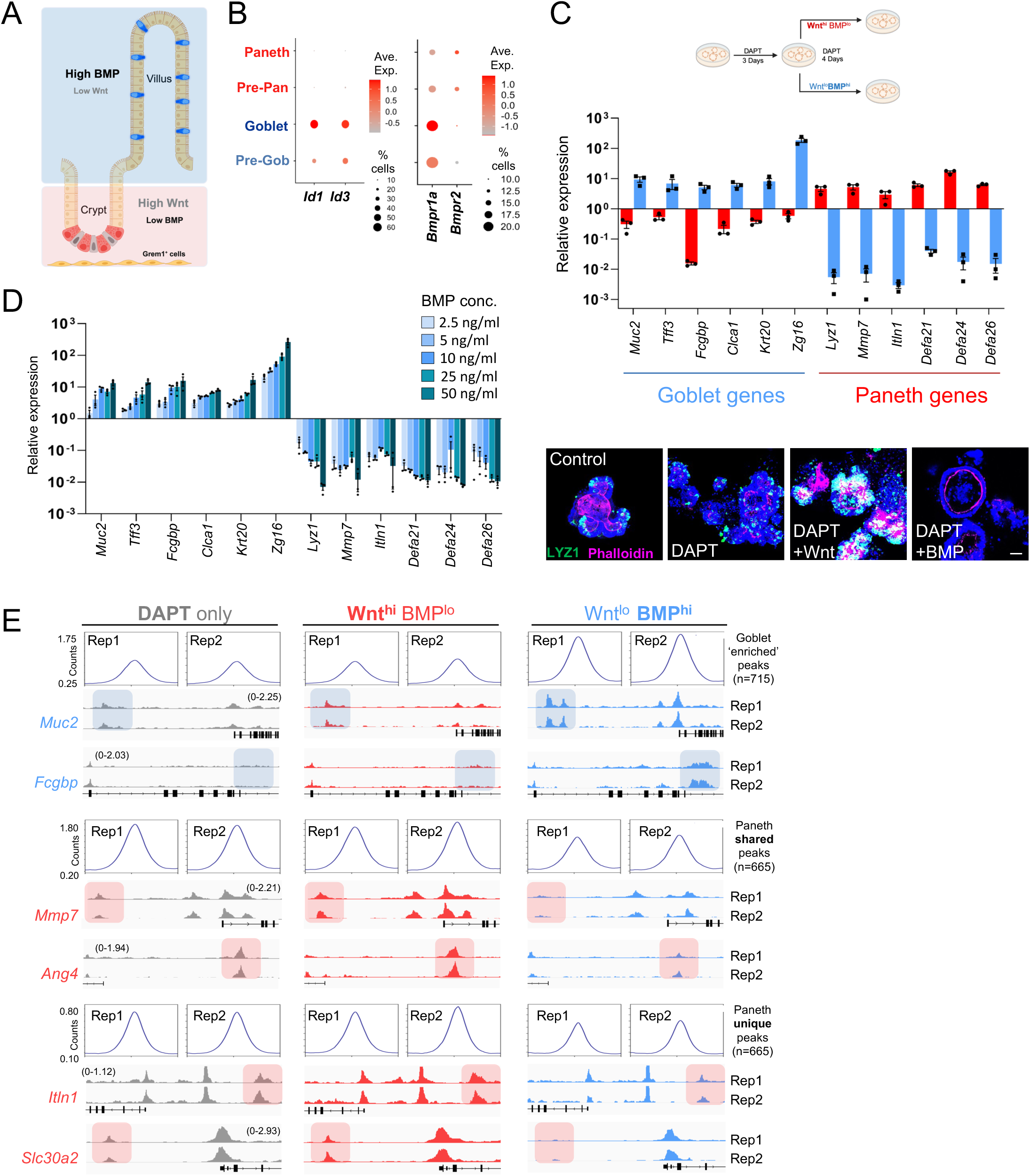
BMP signaling elicits goblet and strongly suppresses Paneth features. **(A)** Sub-epithelial cells expressing various agonists and antagonists generate opposing Wnt and BMP gradients along the crypt–villus axis. **(B)** scRNA data show *Id1* and *Id3* enrichment in goblet cells and BMP receptor gene expression in both Paneth and goblet cells. Circle diameters: within-cluster probability of RNA detection, fill shade: normalized mean RNA level. **(C)** Mouse duodenal organoids were treated with DAPT for 3 days to increase the secretory cell fraction, then cultured under Wnt^hi^ (+CHIR 99021)/BMP^lo^ (+Noggin) or Wnt^lo^ (RSPO1 excluded)/ BMP^hi^ (+BMPs2/4/7, no Noggin) conditions for 4 additional days. RT-qPCR showed 5-to 10-fold increase in Paneth and goblet markers in Wnt^hi^ or BMP^hi^ conditions, respectively, and a massive decline in Paneth markers in the presence of BMP agonists. LYZ1 immunostaining corroborated the findings. n=3 independent experiments, scale bar 50 µm. **(D)** Mouse duodenal organoids were treated with DAPT for 3 days, then in Wnt^lo^ BMP^hi^ conditions with increasing concentrations of the rBMP cocktail. Within a range of 2.5 to 50 ng/mL, RT-qPCR revealed 5-to 15-fold induction of most goblet markers, 20-to 200-fold induction of *Zg16*, and 10-to 200-fold repression of Paneth markers. Each marker gene showed a distinct dose response. n=3 independent experiments. **(E)** Mouse duodenal organoids were treated with DAPT for 3 days, then cultured in Wnt^hi^ BMP^lo^ or Wnt^lo^ BMP^hi^ conditions for 4 additional days and cells were collected for bulk ATAC-seq. In independent replicates, Rep, n=2 each), chromatin accessibility at goblet- and Paneth-enriched sites responded reciprocally to Wnt^lo^ BMP^hi^ conditions. Representative IGV tracks at loci with goblet, Paneth-shared, and Paneth-unique *cis*-elements illustrate increased access at goblet-enriched enhancers and markedly reduced access at both shared and unique Paneth sites in response to BMP signaling. Wnt^hi^ BMP^lo^ conditions had quantitatively small effects on chromatin accessibility. See also Figure S5.

To examine chromatin responses to these niche signals, we performed bulk ATAC-seq on viable cells isolated from organoids treated with 5 μM CHIR 99021 or 50 ng/mL rBMPs. Whereas Wnt^hi^ BMP^lo^ conditions did not affect access at goblet/Paneth sites, Wnt^lo^ BMP^hi^ conditions markedly increased access at goblet sites, and commensurately diminished access at shared and unique Paneth sites (Figure 4E). In contrast, EE- and tuft-selective sites were not materially affected by Wnt or BMP signaling (Figure S5I).

### Phenotypic conversion of post-mitotic cells in vivo and in vitro

To demonstrate the presumptive role of the mesenchymal niche in driving Paneth and goblet phenotypes *in vivo*, we took advantage of the observation that *Grem1*-expressing trophocytes and other fibroblasts dominate in the sub- and peri-cryptal space.^60,65^ To ablate this niche compartment, we treated *Grem1^DTR^* mice^65^ with *Diphtheria* toxin (DT), which rapidly abrogates requisite ISC support.^60,66^ Goblet (AB^+^) and Paneth (LYZ1^+^) cells were zonated and readily distinguished at rest, but 72 h after DT exposure, crypt bases showed dual-stained AB^+^ LYZ1^+^ cells, and by the next day nearly all basal cells lacked LYZ1 and stained with AB (Figure 5A). The speed of alteration, with transient co-staining, indicates that the intact mesenchymal niche actively preserves Paneth –at the expense of goblet– features in crypt base cells.

**Figure 5.**
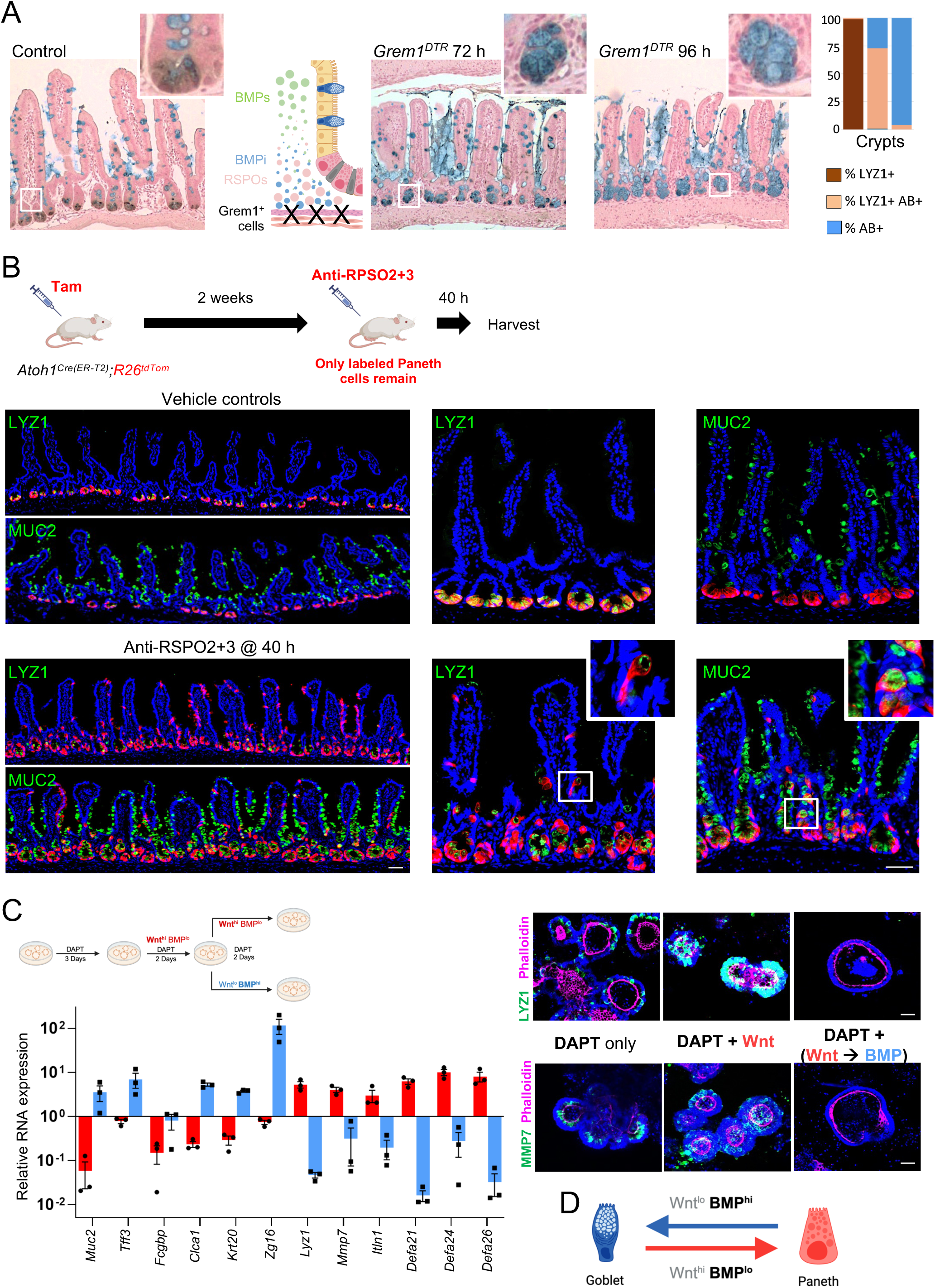
BMP^hi^ and Wnt^lo^ conditions induce a phenotypic switch from mature Paneth to goblet cells *in vivo* and *in vitro*. **(A)** Diphtheria toxin-induced ablation of mesenchymal *Grem1^+^* cells, which is known to remove ISC support, also increased alcian blue (AB) avidity in epithelial LYZ1^+^ cells at the crypt base within 72 h, with LYZ1 loss and AB staining further increased throughout crypts after 96 h, as quantified in the graph. n=4 independent experiments, scale bar 50 µm. **(B)** To capture the phenotypic switch from mature Paneth to goblet cell in response to attenuated Wnt signaling, we marked Paneth cells by injecting Tam into *Atoh1^Cre-ER(T2)^;R26^Tom^* mice, followed 14 days later by IP injection of vehicle or anti-RSPO2 + antiRSPO3 (Anti-RSPO2+3, 10mg/kg) antibodies (Ab). Within 40 h of the latter treatment, labelled (Tom^+^) mature Paneth cells had reduced LYZ1 expression, migrated out of the crypt base, acquired classic goblet morphology (insets), and elevated MUC2 expression compared to vehicle-treated controls, where LYZ1^+^ cells retained Paneth morphology and their location in crypt bottoms. **(C)** Mouse duodenal organoids were treated with DAPT for 3 days, followed by Wnt^hi^ BMP^lo^ conditions for 2 days, then continued in the same conditions or switched to Wnt^lo^ BMP^hi^ conditions (Wnt → BMP) for 2 additional days. RT-qPCR analysis revealed marked attenuation of Wnt-induced Paneth markers and induction of goblet markers under Wnt → BMP conditions. Right: LYZ1 and MMP7 immunostaining corroborate reversal of the Paneth phenotype by BMP signals. n=3 independent experiments, scale bars 50 µm. **(D)** Schema depicting the phenotypic plasticity of post-mitotic goblet and Paneth cells in response to niche signals. See also Figure S5.

In *Grem1^DTR^* intestines, BMP signaling would rise in tandem with attenuated Wnt signaling.^65,67^ To separate the Wnt contribution from that of BMP signaling, we first treated *Atoh1^Cre^*;*R26^Tom^* mice with Tam to label Paneth cells and waited 2 weeks for short-lived goblet and other secretory cells to clear the tissue, while labeled post-mitotic Paneth cells would persist in crypt bases. We then disrupted Wnt signaling by injecting the mice with RSPO2 and RSPO3 antibodies (Ab), which together precipitate loss of Wnt-dependent ISCs.^68^ Within 40 h of Ab treatment, we observed that throughout the small intestine Tom^+^ Paneth cells had reduced LYZ1 expression and a substantial fraction had moved upward, now displaying classic goblet morphology and high MUC2 expression (Figure 5B). This constellation of findings suggests that Wnt signaling retains Paneth cells in the crypt base and promotes Paneth properties; however, even at short distances from crypt bottoms, the Wnt-deficient milieu confers the goblet phenotype on post-mitotic *Atoh1*-marked secretory cells previously showing Paneth features.

To further distinguish reversible signal-mediated goblet/Paneth duality from distinct cell fate assignments, we exposed organoids first to CHIR 99021 and, following washout of that treatment, to rBMP2/4/7. In response to BMPs, cells quickly shed initial Wnt-induced Paneth features and acquired goblet markers (Figure 5C). Together, these findings *in vivo* and *in vitro* reveal Paneth and goblet phenotypes as the composite, interconvertible effects of Wnt^hi^ BMP^lo^ and Wnt^lo^ BMP^hi^ signaling environments, respectively, on labile *Atoh1*-dependent secretory cells (Figure 5D).

### Signal-dependent plasticity of human secretory cells

We asked next if signal-induced phenotypic plasticity underlies human goblet/Paneth duality. Because secretory cell numbers are limiting in human intestinal organoids and mature Paneth cells are especially sparse,^69^ we used 2D cultures of human ISCs (hISCs – previously used to study EE differentiation),^70^ into which we inserted a stable doxycycline (Dox)-inducible *ATOH1-mCherry* (mCh) fusion construct. Exposure to 4 ug/mL Dox for 2 days elevated goblet cell transcripts >100-fold and the two human α-defensins *DEFA5* and *DEFA6* >10^5^-fold (Figure S6A); RNA-seq analysis of EPCAM^+^ cells showed reduced levels of ∼400 ISC transcripts (e.g., *OLFM4*, *GPRC5B*) and robust induction of goblet/Paneth genes (Figures 6A and S6B). Genes induced >2^5^-fold correlated with scRNA profiles^33^ of human goblet and Paneth cells (Figure 6A and Table S3) and those activated 2^2^-to 2^5^-fold included tuft- and EE-specific genes (Figure S6C) such as *SPIB*, *NEUROG3*, and *CHGA*. Thus, hISC^ATOH^^1(Dox)^ constitutes a reliable model to assess human secretory cell responses to Wnt and BMP signaling.

**Figure 6.**
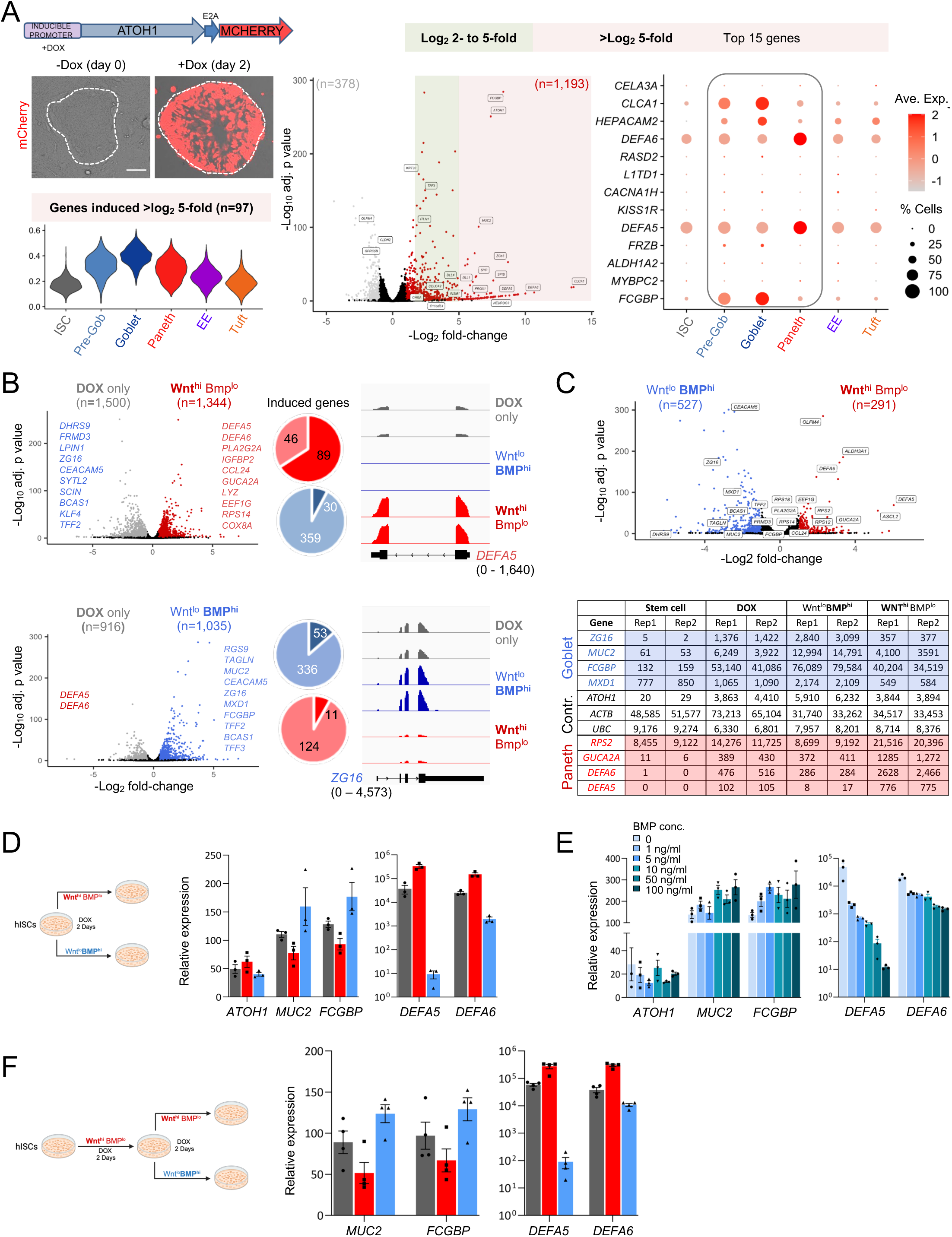
Secretory lineage differentiation from primary human intestinal stem cells *in vitro*. **(A)** A doxycycline (Dox)-inducible *Atoh1-E2A-mCherry* construct was introduced stably into 2D human ISCs to express ATOH1 and mCherry. After 48 h with or without Dox treatment, epithelial (EPCAM^+^) cells were separated from mouse embryonic feeders by FACS and collected for bulk RNA-seq (n=2). Dox exposure induced secretory genes, principally goblet and Paneth markers (>log_2_ 5-fold) but also EE and tuft cell markers (>log_2_ 2.5-fold). Violin plot: representation of genes induced >log_2_ 5-fold in cell types identified in human intestinal scRNA-seq.^33^ Right: Relative expression of the top 15 Dox-induced genes in human scRNA-seq populations. **(B)** Genes differentially expressed (q <0.01, log_2_ fold-difference >0.50) between cells cultured in Dox + Wnt^hi^ BMP^lo^ medium or Dox + Wnt^lo^ BMP^hi^ medium, compared to cells cultured with Dox only (n=2 each, significance of differences determined by the Wald test in DESeq2 default settings). Highly differential genes are named. Pie charts depict the fractions of Paneth (dark red) and goblet (dark blue) genes (from human scRNA-seq data,^33^ increased in response to each treatment. IGV tracks from RNA-seq illustrate representative transcript responses (parentheses: range of signal values). **(C)** Genes differentially expressed (q <0.01, log_2_ fold-difference >1) between cells cultured in Dox + Wnt^lo^ BMP^hi^ and cells cultured in Dox + Wnt^hi^ BMP^lo^ conditions. Significance of differences determined by DESeq2 default settings (Wald test). Table: normalized counts of representative goblet (blue), Paneth (red), and housekeeping (Contr.) genes in untreated and treated conditions. **(D)** RT-qPCR confirmation of selected gene responses to Wnt^lo^ BMP^hi^ (blue bars) and Wnt^hi^ BMP^lo^ (red bars) conditions. Values are normalized to untreated (no Dox) cells. Note different y-axis scales for goblet (left) and Paneth (right) genes. n=3 independent experiments. **(E)** RT-qPCR confirmation of selected gene responses to increasing BMP concentrations. Values are normalized to untreated (no Dox) cells. Note different y-axis scales for goblet (left) and Paneth (right) genes. *DEFA5* was highly sensitive to BMP dosage, *DEFA6* less so, and goblet gene responses varied little in response to BMP doses from 1 ng/mL to 100 ng/mL. n=3 independent experiments. **(F)** After 2 days of culture in Dox + Wnt^hi^ BMP^lo^ conditions, cells were continued in the same condition or switched to Wnt^lo^ BMP^hi^ condition for 2 additional days. RT-qPCR revealed marked attenuation of Wnt-induced Paneth transcripts and modest induction of goblet genes under Wnt → BMP conditions. n=4 independent experiments. See also Figure S6.

To that end, we treated hISC^ATOH^^1(Dox)^ for 2 days with Dox only, with Dox and rBMPs (without RSPO2 or BMP inhibitor DMH-1), or with Dox and CHIR 99021. Bulk RNA-seq analysis showed concordance between duplicate samples and differences resulting from each treatment (Figure S6D and Table S3). Compared to cells treated with Dox alone, Wnt activation in the absence of BMPs induced 1,344 genes, including 89 of the 135 genes (66%, e.g. *DEFA5* and *DEFA6* – Figure 6B) exclusive to human Paneth cells.^33^ In the absence of concomitant Wnt signaling, few of the 1,035 BMP-induced genes were Paneth-enriched and 7.8% (Wnt) to 13.6% (BMP) of goblet-selective genes were altered in opposition to Paneth genes (Figures 6B). Thus, Wnt^hi^ BMP^lo^ conditions favor, and Wnt^lo^ BMP^hi^ conditions potently suppress, Paneth features. Wnt^hi^ BMP^lo^ favored ISC genes such as *ASCL2*, *ALDH3A1* and *OLFM4* (Figure 6C), indicating that both ISC and Paneth are niche-dependent states in humans.

The range of RNA levels between Wnt^hi^ BMP^lo^ and Wnt^lo^ BMP^hi^ conditions was ∼2-to 8-fold for goblet genes but >1 order of magnitude and up to 50-fold (*DEFA5*) for Paneth genes (Figure 6C). RT-qPCR confirmed modest effects on goblet and profound effects on Paneth genes (Figure 6D – note different y-axis scales). Furthermore, while *DEFA6* responded to a dose of 50 ng/mL BMP, with a 5-fold range in response to concentrations between 1 ng/mL and 100 ng/mL, the threshold for *DEFA5* was >50-fold lower and the range was 2,000-fold (Figure 6E). Crucially, Wnt-induced activation of Paneth and suppression of goblet genes were rapidly reversed upon Wnt withdrawal and BMP exposure, with *DEFA5* dropping >3,000-fold and *DEFA6* ∼30-fold (Figure 6F). Thus, human goblet/Paneth duality is niche-mediated, as in mice. Again, highly variable gene responses indicate that local cues do not *specify* Paneth cells but act on individual genes to produce a reversible composite state that has been mistaken for a distinct cell type.

### Cis-regulatory dynamics during intestinal epithelium differentiation

Our observations establish a framework to consider *cis*-element dynamics not only in the secretory lineage but broadly for unitarian ISCs. Having mapped accessible chromatin in *Atoh1*-labeled ISCs by snATAC-seq (Figure 2E) and in purified duodenal Lgr5^+^ ISCs by bulk ATAC-seq (Figure S3H), we compared the latter data with new bulk ATAC-seq data on duodenal enterocytes purified from *Vil1^CreER-T^*^2^*;Atoh1^Fl/Fl^* mouse villi, which lack all secretory cells.^10,11^ In line with prior reports,^12,71,72^ most enterocyte enhancers were comparably open in ISCs; only 21.73% of sites (n=9,429) showed less or more accessibility in one population or the other (log_2_ fold-difference ≥1.5, FDR ≤0.001, Figure 7A and Table S4), affirming that enterocyte enhancer access is largely pre-determined in ISCs. In the bulk ATAC-seq data from ISCs and purified enterocytes, we then assessed the enhancers enriched in individual secretory cell types (snATAC-seq, Figure 2E). For this analysis, we excluded promoter regions and included 4,684 enhancers enriched in *Atoh1*-labeled crypt enterocyte progenitors; these steps refined the numbers of ISC- and secretory-specific sites we had considered earlier but the number of goblet-enriched enhancers remained small (Figure 7B vs. Figure 2E). While EE-, tuft-, and Paneth-unique sites were weakly accessible in ISCs, goblet-enriched and especially Paneth-shared enhancers were at least as accessible as the enhancers enriched in enterocyte progenitors (Figure 7B and Table S5). Thus, specific TFs activate large sets of unique enhancers to establish EE or tuft cell identities, but goblet/Paneth features arise largely through sites that, like enterocyte enhancers, are already accessible in ISCs (Figure 7C). Building on prior observations that ISC chromatin is “primed” for enterocyte or goblet/Paneth differentiation,^12^ these findings provide mechanistic parallels to the unitarian model. First, they reveal that while enterocyte enhancers remain open and may become more accessible in mature enterocytes, chromatin accessibility at goblet/Paneth sites declines in those cells (Figures 7B and 7C). Second, they indicate that barring activation of Paneth-unique sites under Wnt^hi^ BMP^lo^ conditions or expression of NEUROG3 or POU2F3 to induce EE or tuft cell fates, the natural outcome of the dominant ATOH1^+^ secretory population is to display goblet properties.

**Figure 7.**
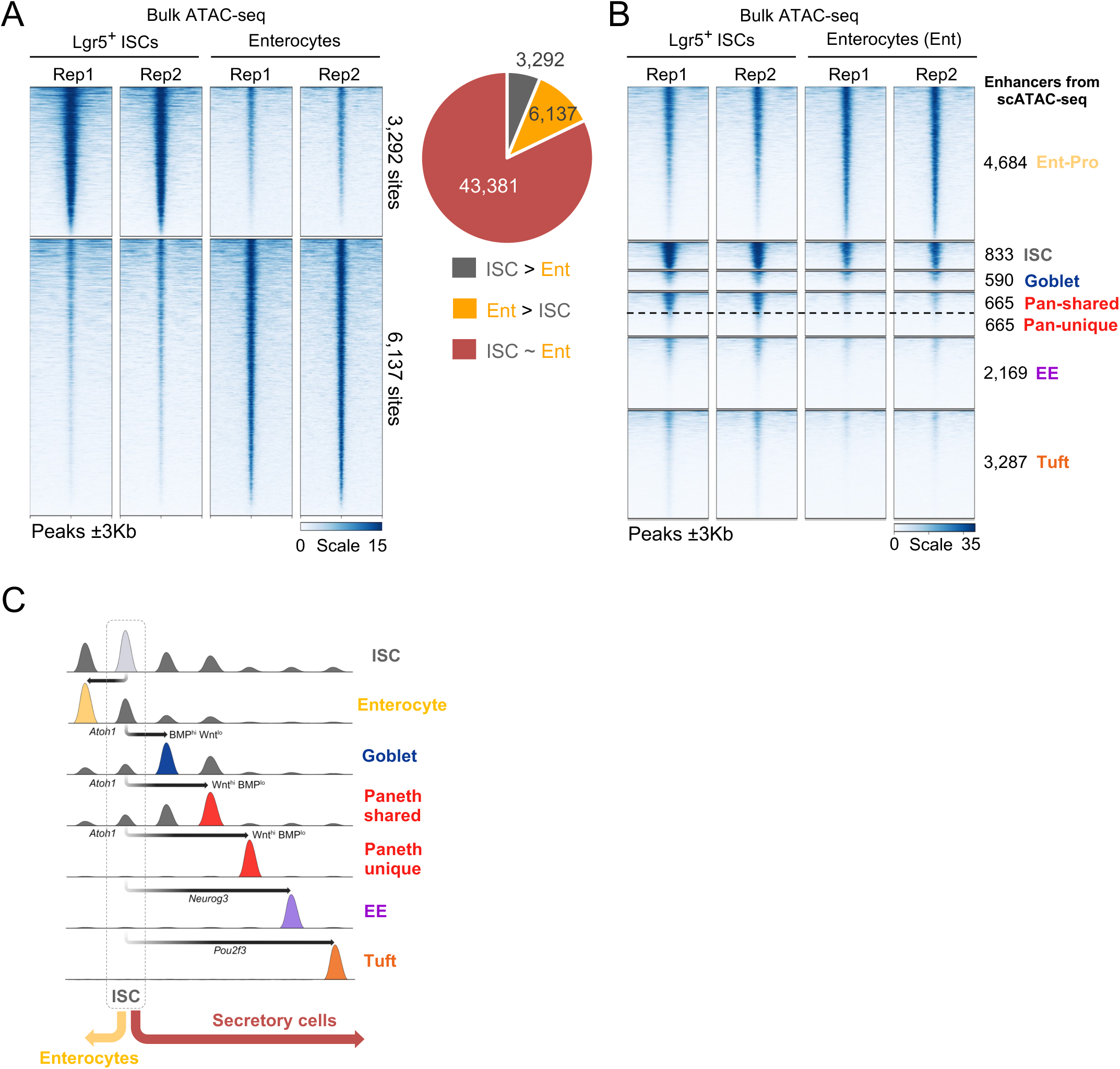
Chromatin accessibility of secretory-cell enhancers in LGR5^+^ ISCs and *Atoh1*-null villus enterocytes. **(A)** Most of the 52,805 enhancers identified in bulk ATAC-seq of duodenal ISCs and villus *Atoh1^-/-^* enterocytes (Ent) were equally accessible in both. Only 3,292 enhancers and 6,132 enhancers were more accessible (log_2_ fold-difference ≥1.5, FDR ≤0.001) in ISCs or Ent, respectively, confirming prior reports that the bulk of the Ent epigenome is programmed in ISCs. n=2 replicates (Rep) of each cell type. **(B)** Status of enhancers that are differentially accessible in each *Atoh1*-labeled cell type in purified duodenal LGR5^+^ ISCs and *Atoh1*-null villus enterocytes (Ent). ISCs share most open enhancers with villus Ent, followed by goblet and Paneth-shared enhancers. In villus Ent, accessibility of Ent-active enhancers is measurably increased relative to ISCs, while the accessibility of goblet-active and Paneth-shared enhancers is reduced. Paneth-unique, EE, and tuft enhancers are largely inaccessible in ISCs and differentiated Ent. **(C)** *Cis*-regulatory dynamics of a revised unitarian model for intestinal epithelium.

## DISCUSSION

Our findings challenge the long-held view that goblet and Paneth cells are distinct terminal ISC derivatives. We show that these cells are not singularly specified, in the manner that restricted TFs enable selective chromatin access at thousands of EE and tuft cell enhancers. Rather, goblet and Paneth properties are alternative and reversible manifestations of a single, versatile *Atoh1*-dependent secretory cell in response to niche-derived Wnt and especially BMP signals. Another previously unrecognized feature is that a few anti-microbial genes account for 1/3 (goblet) to 2/3 (Paneth) of all transcripts. Differential –but non-exclusive– expression of these genes (e.g., *Muc2*, *Tff3*, *α-Defensins*, *Lyz1*) has historically distinguished the two cell types and they are prominent among the genes that respond reciprocally to Wnt^lo^ BMP^hi^ or Wnt^hi^ BMP^lo^ conditions and various microbial exposures. Ectopic Wnt activation induced a goblet-to-Paneth phenotypic switch largely confined to crypts and lower villi. Conversely, RSPO attenuation rapidly induced upward migration of mature, post-mitotic Paneth cells and their acquisition of goblet features. These observations explain why the Paneth phenotype is confined to the Wnt^hi^ BMP^lo^ milieu in crypt bottoms.^67^

Wnt signaling was previously implicated in Paneth cell ‘differentiation,^50,51,53^ but we found that it affected goblet/Paneth signature genes modestly compared to BMPs, which potently suppress Paneth-selective genes. Taken together, our findings implicate Wnt/Rspo signals in retaining ATOH1-specified secretory cells at the crypt base, where the severe paucity of BMP activity helps boost expression of Paneth-restricted genes by orders of magnitude. High BMP signaling from sub-epithelial cells at villus tips^60,73^ likely further limits expression of the Paneth phenotype in healthy villi. Gene responses to BMPs vary considerably, further attesting that goblet and Paneth are not distinct cell types, but composite phenotypes of distinct antimicrobial genes expressed in response to local Wnt and BMP concentrations. These same signals control the ISC state^67^ and, albeit crucial, are not the only influences. MAP kinase signaling downstream of receptor tyrosine kinases and oncogenic *Kras*, for example, affect ISC growth and the balance between Paneth and goblet properties.^74,75^ Additional niche factors likely affect ISCs, secretory cells, or both.

Beyond revising the unitarian model and linking it firmly to instructions from the underlying mesenchyme, our findings identify the underlying chromatin dynamics. *Cis*-elements common to enterocytes and goblet/Paneth cells are accessible even in ISCs, and select modules become accentuated in secretory cells (*Atoh1^+^*) or enterocytes (*Atoh1^-^*) at the relative expense of the other. While goblet and most Paneth sites are shared, ∼650 Paneth-exclusive sites lie alone or interspersed with shared sites at Paneth-expressed loci. Both shared and unique sites respond to Wnt^lo^ BMP^hi^ signaling conditions, and the goblet phenotype appears to be the default state of *Atoh1*^+^ secretory ISC daughters. Most of these daughters depart from the crypt base, where they encounter niches with less Wnt and more BMP signaling and use previously accessible enhancers to robustly activate antimicrobial genes such as *Zg16*, *Tff3*, *Fcgbp*, *Clca1*, and *Muc2*. The minority of *Atoh1*^+^ cells that remain in the Wnt^hi^ BMP^lo^ space deep in the crypt base use other previously accessible enhancers and recruit additional sites unique to the Paneth phenotype to express a different complement of antimicrobial genes, especially *α-Defensins*. In response to unknown signals, other *Atoh1*^+^ cells activate *Neurog3* or *Pou2f3*, which recruit distinct *cis*-elements –directly or indirectly– to specify alternative EE or tuft cell fates. Our findings do not address whether the latter cell types are interconvertible, but *Neurog3^+^* cells can dedifferentiate into ISCs^17^ and tuft cells can arise from *Atoh1*^-^ ISC progeny.^41,42^ One question still unanswered is whether microbe-or inflammation-induced expression of Paneth genes in villus goblet cells occurs because local Wnt/BMP gradients get perturbed, because other niche signals exert similar effects, or by recruiting *cis*-elements that are inaccessible during homeostasis and were therefore not captured in our molecular studies of resting intestines.

## Acknowledgments

Supported by NIH grants R01DK081113 and R01DK082889 (R.A.S.), the Claudia Adams Barr program in basic cancer research, and generous gifts from the Lind family. We thank M. Hoshino (*Atoh1^Cre^*), S. Robine (*Vil1-Cre^ER-T^*^2^), and C. Perret (*Apc^fl^*) for kind gifts of mouse strains; E.E. Storm and F.J. de. Sauvage for the gift of RSPO2 & RSPO3 antibodies; S. Madha, Rong Li, and H. Long for bioinformatic assistance; K. Badarinath for valuable discussions; the Harvard Digestive Disease Center (P30DK034854) for organoid culture reagents; and the staff of flow cytometry and microscopy facilities at the Dana-Farber Cancer Institute.

## Author Contributions

S. Bhattacharya and R.A.S. conceived and designed the study; S. Bhattacharya, G.T., E.M., O.E., and J.K. performed experiments; S. Bhattacharya, P.N.P.S., R.H., and J.K. performed computational analyses; W.G. and Q.Z. provided human ISCs; Y.P. assisted with genotyping and cell quantitation; T.C.W., S. Beyaz, and R.A.S. supervised various parts of the study; S. Bhattacharya and R.A.S. wrote the paper, with input and approval from all authors.

## METHODS

## RESOURCE AVAILABILITY

### Lead Contact

Further information and requests for resources and reagents should be directed to and will be fulfilled by the lead contact, Ramesh Shivdasani.

### Materials availability

Human organoid-derived lines generated in this study are available and can be requested from the lead contact; a Materials Transfer Agreement may be required.

### Data and code availability

Sequencing data have been deposited in the Gene Expression Omnibus (GEO) under series GSE271611, GSE271613, GSE271614, GSE271615.

This paper does not report any original code. Any additional information required to reanalyze the data reported in this paper is available from the lead contact upon request.

## EXPERIMENT MODEL AND STUDY SUBJECT DETAILS

### Animals

*Atoh1^Cre^* mice,^25^ *Vil1-Cre^ER-T^*^2^ mice,^76^ and *Apc^Fl^* mice,^77^ were generous gifts from M. Hoshino (National Center of Neurology and Psychiatry, Japan), S. Robine (Institut Pasteur, France), and C. Perret (Institut Cochin, France), respectively, and *Grem1^DTR^* mice were described previously.^65^ *R26R^Tom^* (JAX strain 007909),^78^ *Lgr5^CreER-T^*^2^ (JAX strain 008875),^3^ and *Atoh1^fllfl^* (JAX strain 008681),^11^ mice were purchased from Jackson Laboratories. All strains were maintained on a predominant C57BL/6 background and housed under specific pathogen-free conditions in 12 h light/dark cycles at 23 ±1°C, 55 ±15% humidity, with *ad libitum* access to food and water. Animals were weaned 21 to 28 days after birth and handled according to procedures approved by Animal Care and Use Committees at the Dana-Farber Cancer Institute, Columbia University Medical Center, or Cold Spring Harbor Laboratory. Experimental mice and littermate controls of both sexes were 8 to 28 weeks old at the time of experiments and cell isolations.

### Cell lines

HEK293FT cells (female fetus, Invitrogen, R70007) were cultured in Dulbecco’s Modified Eagle Medium (DMEM, Gibco, 11965092) containing 10% fetal bovine serum (FBS, Corning, 35-010-CV) and 1x Penicillin/Streptomycin (Gibco, 15140163) at 37°C, 5% CO_2_.

### *Homo sapiens*: hISC^Atoh^^1(Dox)^ primary cells

The process to generate primary human intestinal stem cells (hISCs) was described recently;^70^ hISCs were cultured over feeder layers of mitotically inactivated mouse embryonic fibroblasts and engineered for inducible ATOH1 expression. They were cultured in 90% basal medium (60% high-glucose DMEM (Gibco, 11965092), 20% F12K medium (Gibco, 21-127-022), and 20% FBS (Corning 35-010-CV)) and 10% RSPO2-conditioned medium supplemented with 10 mM nicotinamide (Sigma-Aldrich, N5535), 25 µM Primocin (Invivogen, ant-pm-1), 1 µM A8301 (Sigma-Aldrich, SML0788), 5 µg/mL Insulin (Sigma-Aldrich I0516), 10 µM Y27632 (LC Laboratories Y5301), 1 µM DMH1 (a BMP inhibitor, Sigma-Aldrich D8946), 50 ng/mL epidermal growth factor (EGF, R&D, 236EG200) and 2 µM triiodo L-thyronine (Sigma-Aldrich, T3697) at 37°C in a humidified atmosphere of 7.5% CO_2_.

### *Mus Musculus*: Primary mouse embryonic fibroblasts

DR4 mouse embryonic fibroblasts (MEFs), male and female mixed, were obtained from the American Type Culture Collection (ATCC, SCRC-1045) and cultured in DMEM (Gibco, 11965092) containing 10% FBS and 1x Penicillin/Streptomycin at 37°C in 5% CO_2_.

## METHOD DETAILS

### Mouse treatments

To activate CRE^ER-T^^2^ in *Atoh1^CreER-T^*^2^;*R26R^Tom^* mice, we administered 2 doses of 1 mg/mL tamoxifen (Sigma-Aldrich, T5648; stocks prepared in cornflower oil) by intraperitoneal (IP) injection 24 h apart; we harvested intestines 12 h later for immunostaining, scRNA-seq or scATAC-seq. To delete *Atoh1* in *Atoh1^Cre/Fl^*;*R26^Tom^* mice, we administered 1 mg/ml tamoxifen IP once and harvested intestines after 3 or 10 days. To delete *Apc* in *Atoh1^Cre^*;*Apc^fl/fl^*; *R26R^Tom^* mice for scRNA-seq or snATAC-seq (Figure 3), we injected 2 doses of 1 mg/ml tamoxifen IP 24 h apart and harvested intestines 12 h later. For immunostaining (Figures 3 and S5), these mice received single IP injections of 1 mg/ml tamoxifen IP and 10 μM of 5-ethynyl-2′-deoxyuridine (EdU) and we harvested intestines 24 h later. Anti-RSPO2 and Anti-RSPO3 antibodies^68^ gifted by F. de Sauvage (Genentech) were injected IP together at doses of 10 mg/kg each, and intestines were harvested 40 h later.

For *Strongyloides venezuelensis* infection (Figure 2), mice were injected subcutaneously with 500 L3 larvae, left undisturbed for 14 days, and euthanised for intestinal harvest on day 15. For *Salmonella typhimurium* (*Stm*) infection (Figure 2), we used strain IR715 (Ref. ^79^) a nalidixic acid-resistant derivative of *Stm* strain 14028 (American Type Culture Collection). Mice were pre-treated with streptomycin (Fisher Scientific, BP910-50, 0.2 g/mL in phosphate-buffered saline (PBS)) 1 day before intragastric inoculation with 0.1 mL PBS or 10^8^ CFU of *Stm* suspension (inoculum titer confirmed by plating 10-fold serial dilutions on LB agar plates containing 50 mg/mL nalidixic acid, Alfa Aesar, J63550-06); food was removed for 4-6 h prior to inoculation and resumed *ad libitum* afterward. Intestines harvested 5 days later were fixed in 10% formalin.

Age- and sex-matched C57BL/6J mice received intraperitoneal injections of IL-25 (Figure 2). (R&D Systems, 1399-IL-025, 500 ng) in 200 µl PBS daily for 7 days and intestines were harvested on the 8^th^ day. For succinate treatment, mice were provided with regular drinking water or 150 mM sodium succinate dibasic hexahydrate (Sigma-Aldrich, S2378) *ad libitum* for 7 days and intestines were harvested on the 8^th^ day.

Mice received a single dose of adenovirus carrying RSPO1-FC or Control-FC (Figure 3, details about virus production and administration were described previously)^58^ and intestines were harvested at the indicated times. *Grem1^DTR^* mice (Figure 4) were treated with two IP injections of Diphtheria toxin (Sigma-Aldrich, D0564, 50 μg/kg) on alternate days. *Vil1-Cre^(ER-T^*^2^*^)^;Apc^fl/fl^* mice (Figure S5) were injected with 2 IP doses of 1 mg/ml tamoxifen 24 h apart and intestines were harvested 7 days later. To minimize residual long-lived Paneth cells, *Vil1^CreERT^*^2^*;Atoh1^fl/fl^* (*Atoh1*-null) mice (Figures 7) were sampled 3 weeks after treatment with 1 mg/mL tamoxifen.

### Detection of tdTom^+^ fluorescent cells in tissues

Mouse small intestines were fixed in 4% paraformaldehyde (Electron Microscopy Sciences, 15714-S) in phosphate-buffered saline (PBS) at 4°C overnight with gentle agitation, washed 3 times in PBS at 4°C, incubated through a 10%– 30% sucrose gradient, embedded in OCT compound (Tissue-Tek, 4583), and stored at −80°C. Tissue sections (8 μm) adhered onto glass slides were washed in PBS, mounted in medium containing 4′,6-diamidino-2-phenyl-indole (DAPI, Vector Laboratories, H-1200), and fluorescent cells were visualized and counted using a Zeiss LSM 980 confocal microscope. Images were analyzed by Fiji software.^80^

### Intestinal crypt and villus isolations

Pieces of small intestine were rinsed in PBS, then rotated in 5 mM EDTA (pH 8) in for 45 min at 4°C with manual shaking every 10 min and a change of solution after 30 min. Released epithelium was passed through a 70-µm strainer to separate villi from crypts and each fraction was dissociated into single cells by rotating in 4% TrypLE solution (ThermoFisher, A1217702) for 30-45 min at 37°C. Single cells were diluted in Dulbecco’s modified Eagle medium (DMEM, Corning. 17-205-CV) containing 2% FBS. Tom^+^ cells were isolated on a FACSAria II SORP flow cytometer, with elimination of dead (DAPI^+^) cells, pelleted by centrifugation, and trypan blue-excluding cells were counted manually.

### Fluorescence-activated cell sorting

Viable (DAPI^-^) Tom^+^ mouse cells (Figures 1 and 2) and EPCAM^+^ hISCs (Figure 6) were sorted using a BD FACSAria II instrument (BD Biosciences) with a 70 µm nozzle. Cells were stained for EPCAM (Invitrogen, 46-5791-82, 1:100) for 30 min at 4°C in Hank’s Balanced Salt Solution (HBSS, Gibco, 14065-056) containing 10 mM HEPES, 2 mM EDTA, and 0.5% fetal bovine serum albumin (FBS, Corning, 35-010-CV). hISCs were collected in HBSS supplemented with 0.9% glucose, 10 mM HEPES, 10 µM Y27632, and 2% FBS, and washed in PBS. Data were analyzed in FlowJo software (BD Life Sciences).

### Single-cell RNA sequencing

8,000-10,000 DAPI^-^ cells, sorted by flow cytometry, were loaded onto a 10X Chromium controller followed by the Chromium Next GEM Single Cell 3′ V3.1 assay (10X Genomics, PN-1000121). Libraries were constructed according to the manufacturer’s recommendations and sequenced on the NovaSeq platform (Illumina).

### Single-nucleus ATAC sequencing

8,000-15,000 DAPI^-^ cells, sorted by flow cytometry, were loaded onto a 10X Chromium controller followed by the Chromium Next GEM Single Cell ATAC Reagent Kits v1.1 (10X Genomics, PN-1000175). Libraries were constructed according to the manufacturer’s recommendations and sequenced on the NovaSeq platform (Illumina).

### Bulk RNA-sequencing

5×10^4^ viable cells (DAPI^-^ by flow cytometry) were used immediately for RNA extraction with RNAeasy Plus micro kit (Qiagen). RNA integrity and concentration were determined using Agilent Bioanalyzer High Sensitivity RNA 6000 pico kits (Agilent). For samples with RNA-integrity number (RIN) >8.5, libraries were constructed using SMART-Seq v4 Ultra Low Input RNA kits and sequenced on the NovaSeq platform (Illumina).

### Bulk ATAC-sequencing

5×10^4^ DAPI^-^ cells, sorted by flow cytometry were used for ATAC-seq as described previously.^81^ Cells were washed in ice-cold PBS and lysed in 50 µL ice-cold ATAC resuspension buffer (10 mM Tris-HCl pH 7.4, 10 mM NaCl, 3 mM MgCl2, 0.1% IGEPAL CA630, 0.1% Tween-20, 0.01% digitonin). Lysed nuclei were washed in 1 mL ATAC resuspension buffer without IGEPAL or digitonin and centrifuged at 500 *g* for 10 min at 4°C. Nuclear pellets were treated with 50 µl Nextera Tn5 Transposase mixture (Illumina, 20034197) for 30 min at 37°C.

Transposed chromatin was purified with QIAquick PCR purification kits (Qiagen, 28004) and preamplified with high-fidelity 2X PCR master mix (New England Biolabs) using a universal forward primer and unique reverse primers in a 50 µl reaction for 5 cycles. 5 µl of these pre-amplified fragments were subjected to 20 cycles of qPCR amplification to determine additional cycles (1/3 of the maximum qPCR fluorescence intensity) for the remaining 45 µl. After the final amplification, libraries were purified using Agencourt AMPure XP beads (Beckman Coulter, A63880) to remove primer dimers. Library quality and size distribution were analyzed using Agilent High Sensitivity DNA Bioanalysis chip (Agilent, 5067-4626). Libraries were sequenced on a NovaSeq 6000 instrument (Illumina) to generate 150-bp reads.

### Immunofluorescence, immunohistochemistry, and whole-mount staining

Intestines were harvested at the specified times, rinsed in PBS, and fixed overnight in 4% paraformaldehyde. For immunohistochemistry, fixed tissues were incubated through a 10%–30% sucrose gradient, embedded in OCT compound (Tissue-Tek, VWR Scientific 4583), and 8-µm sections were prepared using a Leica cryostat. After blocking for 1 h with 0.5% BSA, slides were incubated with antibodies (Ab) against CHGA (Abcam, ab15160, 1:100), LYZ1 (Dako, A0099, 1:1000), MUC2 (Santa Cruz Biotechnology, sc15334, 1:1000), DCLK1 (Abcam, ab31704, 1:200) or MMP7 (Cell Signaling Technology, S3018, 1:250), at 4°C overnight, followed by washes and incubation with donkey anti-Rabbit IgG conjugated with Alexa Fluor 488 or Alexa Fluor 647 (Invitrogen). To determine S-phase fractions, EdU^+^ cells were stained using Click-iT^TM^ EdU Cell Proliferation kit (Invitrogen, C10640) according to the manufacturer’s recommendations. After counterstaining with DAPI, slides were mounted with Vectashield (Vector Laboratories), imaged on a Zeiss LSM 980 confocal or Leica Thunder Imager microscope, and processed using Fiji software.^80^

Goblet cells were visualized using the Alcian Blue Stain Kit (Figures 3 and 5) (pH 2.5, Mucin Stain, Abcam, ab150662) according to the manufacturer’s recommendations. To visualize Paneth cells, slides were incubated with LYZ1 Ab (1:1000, Dako, A0099) at 4°C overnight, secondary goat anti-rabbit IgG (H+L, Vector Laboratories, BA-1000-1.5, 1:400) for 1 h, and a chromogenic substrate (ImmPACT® DAB Substrate Kit, Vector Laboratories, SK-4105) before mounting in Vectashield (Vector Laboratories).

After removal of Matrigel with cell recovery solution (Corning 354253), mouse duodenal organoids were fixed in 4% paraformaldehyde for 4 h, blocked with 0.2% bovine serum albumin and 0.1% Triton X-100 (blocking solution) for 30 min, and stained with LYZ1 (Dako, A0099, 1:1000) or MMP7 (Cell Signaling Technology, S3018, 1:250) overnight at 4°C. Organoids were washed with blocking solution, incubated overnight with appropriate secondary Ab (Alexa fluor, Invitrogen), DAPI, and phalloidin (Life Technologies, A22287, 1:500), washed again with blocking solution, mounted with spacer (Grace Bio-Labs, 654002) and Vectashield (Vector Laboratories, H-1000). To determine S-phase fractions (Figure S5), cells were labeled with 10 μM EdU for 30 min and stained using Click-iT EdU Imaging Kit, Alexa Fluor 647 dye according to the manufacturer’s recommended protocol. Images were taken on a Zeiss LSM 980 confocal microscope. Images were processed using Fiji software.^80^

### Organoid cultures

Between 50 and 100 crypts isolated from the above procedure were added to 10 μl or 20 μl Matrigel (Corning) droplets, respectively. Organoid culture medium (ENR), as described previously,^62^ contained Advanced DMEM, 1% B27 (Gibco), 0.5% N2-supplement (Gibco), 125 mM N-Acetylcysteine (Sigma), 10% Rspo1-conditioned medium (Harvard Digestive Disease Center), 100 ng/ml rNOG (Peprotech), 100 ng/ml rEGF (Peprotech) and 10 μM Y-27632 (Sigma). To induce secretory differentiation (Figures 4 and S5) organoids were treated with DAPT (10 μM, Sigma-Aldrich D5942) for 3 days and then continued on DAPT for 4 additional days as control or treated with DAPT and GSK3-inhibitor CHIR99021 (Sigma, 5 μM) to mimic canonical Wnt-activation in Wnt^hi^BMP^lo^ conditions or with DAPT and mixtures of recombinant(r) BMPs (rBMP2/4/7 combined, 50 ng/ml each) without rNOG and RSPO1 for Wnt^lo^ BMP^hi^ conditions for 4 additional days.

### RNA extraction, reverse transcription, and RT-qPCR

Freshly sorted DAPI^-^ cells were pelleted and lysed in RLT Plus buffer (Qiagen) and RNA was extracted using RNeasy Plus Micro kit (Qiagen). Cultured cells or isolated intestinal crypts were lysed in TRIZOL reagent (Life Technologies) and RNA was extracted by combining TRIZOL-chloroform extraction with RNA-binding columns (PureLink kit). RNA (1 μg) was reverse transcribed using SuperScript kits (Life Technologies) and transcripts were quantified on a BioRad instrument using Power SYBR reagents (Life Technologies).

### Generation of hISCs with inducible ATOH1 activity

Human *ATOH1* was amplified from Addgene plasmid 162342 (Ref. ^82^) and used to replace mouse *Neurog3* in the vector pLenti-rtTA-Blast^R^-TetO-*Neurog3-mcherry* (Ref. ^83^) by enzymatic assembly.^84^ Lentiviral particles (10^8^ transduction units/mL) were produced using HEK293FT cells (Invitrogen, R70007) as described previously.^83^ hISC cultures were transduced with lentiviral particles carrying PGK-rtTA-Blast^R^ and TetO-*ATOH1-mcherry* in the presence of 10 µg/mL polybrene. After 48 h, cells were selected in 10 µg/mL blasticidin for 7 days. hISC^ATOH^^1(Dox)^ cells were induced with 4 µg/mL doxycycline (Sigma-Aldrich, D9891) and ATOH1 expression was verified by visualizing mCherry fluorescence. To examine ATOH1 effects, engineered hISCs were separated from MEF feeder cells by flow cytometry for EPCAM.

## DATA PROCESSING AND ANALYSIS

### scRNA sequencing

Fastq files were aligned to mouse genome version mm10, unique molecular identifiers (UMIs) were counted using Cell Ranger (10X Genomics) v3.0.2 with default parameters, and quality control, normalization, and clustering were performed in Seurat package v4.3.0.1 (Ref. ^85^) in R version 4.2.0 (https://www.r-project.org/). Cells with >900 and <3,000 features in duplicate libraries from *Atoh1^Cre^*;*R26^Tom^ (wild-type, WT)* (Figure S1) or *Atoh1^Cre^*;*Apc^fl/fl^* (*Apc^-/-^*) duodenum (Figure 3), with >600 and <3,000 features in duplicate libraries from *WT* Ileum (Figure S1), and <7% mitochondrial genes in all samples were retained. These filters yielded 8,693 cells from *WT* duodenum, 6,723 cells from *Apc^-/-^* duodenum, and 11,598 cells from *WT* ileum. *WT* and *Apc^-/-^* duodenal data were merged and analyzed using the ‘NormalizeData’ function, whereas *WT* duodenal and *WT* ileal data were merged and integrated using ‘FindIntegrationAnchors’ and ‘IntegrateData’ functions in Seurat. Cell features were detected with the ‘FindVariableFeatures’ function (2,000 features, selection method “vst”) and unwanted sources of variation, such as mitochondrial gene expression, were regressed out. Principal component analysis (PCA) was performed on the scaled data using ‘RunPCA’. Dimensionality was reduced using Uniform Manifold Approximation and Projection (UMAP) on the first 21 dimensions to place cells in similar local neighborhoods close together in two-dimensional space and K-nearest neighbor (KNN) graphs were constructed using the ‘FindNeighbors’ function. We clustered cells using the ‘FindClusters’ function, which implements an optimal Louvain algorithm to group cells together iteratively by nearest-neighbor modules and we used the ‘FindAllMarkers’ function with default parameters to identify genes enriched in each cluster, which was assigned a cell identity based on the profile of highly enriched genes (Supplemental Figure S1D).

### Cell type similarity

We assessed similarities among duodenal and ileal cell types (Figure 1) using MetaNeighbor analysis.^86^ Highly variable genes were used to construct a cell-cell similarity network and a neighbour-voting algorithm was then used to predict cell type identities of individual cells given their neighbours. The area under the receiver operating characteristic (AUROC score) of this prediction was used as a metric of similarity.

### Comparisons with published datasets

scRNA-seq data from control and infected mouse intestinal epithelium^32^ (GEO: GSE92332, Figure S2), Tom^+^ cells from *Neurog3^Cre-ER(T^*^2^*^);^R26^Tom^* mouse intestinal epithelium^17^ (GEO: GSE183299, Figures S4), and human jejunal epithelium^33^ (HuBMAP: HBM692.JRZB.356, https://doi.org/10.5061/dryad, Figures 1, 6, S2, S3, S4 and S5) were used to validate authenticity of prominent genes and cell states identified in mouse and human (hISC^Atoh^^1(Dox)^) secretory cells. Cells from published datasets were filtered for stem and secretory cell clusters, and then renormalized before further analysis.

### Bulk RNA sequencing

Libraries were aligned to mouse genome version mm10 or human genome version hg19 and processed using Viper (which utilizes STAR aligner^87^) with default parameters to generate raw counts.^88^ Raw counts were normalized and differential gene expression was determined using DESeq2 v1.36.0 (Ref. ^89^) in R software v4.2.0 (https://www.r-project.org/). Plots were generated using the ggplot2 v3.3.5 package^90^ in R or EnhancedVolcano plot (https://github.com/kevinblighe/EnhancedVolcano).

### Bulk ATAC sequencing

Sequences were aligned to mouse genome version mm10 using bowtie2 v2.3.4.3 with default parameters.^91^ Peaks were called using MACS2 v2.1.1,96 (Ref. ^92^) with q-value cut-off 0.001, lower Mfold=5, upper Mfold=50. ATAC peaks from duplicate samples were concatenated and merged for overlapping regions using BEDTools.^93^ Bigwig files were generated and normalized using haystack v0.5.5 (Ref. ^94^), followed by deep tools2 v3.4.3 (Ref. ^95^) to plot heatmaps (descending order of signal strength at peak summits ±3 kb across each cluster) and to visualize read distributions on Integrative Genomics Viewer v2.5.0 (Ref. ^96^).

### snATAC sequencing

Fastq files were aligned to mouse genome version mm10. Transposase cut sites and peak accessibility were identified using Cell Ranger ATAC pipeline v2.0.0 (https://github.com/10XGenomics/cellranger). Data were analyzed using the Signac package v1.7.0 in R version 4.1.1 (https://github.com/timoast/signac). We retained cells with >2,500 and <30,000 peak fragments, >30% reads within peaks, a genome blacklist ratio <0.075, nucleosome siganal <4 and TSS (Transcription Start Site) enrichment score >2. These filters yielded 15,145 cells (Figure 2) in the *WT* control duodenum (n=2). The same filters yielded 26,331 cells (Figures 3 and S5) in *APC^-/-^* duodenum (n=3). Data were normalized using the Seurat extension Signac,^97^ by running term frequency-inverse document frequency (TF-IDF) and singular value decomposition (SVD) on the TF-IDF normalized matrix to reduce dimensionality. Non-linear reduction was performed with UMAP and graph-based clustering with the Louvain algorithm. Gene activity scores were computed using chromatin accessibility reads associated with each gene. scRNA-seq cell clusters corresponding to scATAC-seq clusters were predicted using FindTransferAnchors and TransferData functions (Figure S3). Once clusters were annotated, peaks were identified using the ‘CallPeaks’ function in MACS2 (Ref. ^92^) with default parameters. Peaks overlapping with annotated blacklist_v2 regions in mm10 were removed. To access the fraction of overlapping peaks, EDLC peaks were excluded owing to their low cell number. Disregarding peaks called in every other cluster, overlapping peak fractions (Figure 2) were defined as the ratio of peaks shared by two cell types to the total number in the cell type with fewer peaks. ATAC sites enriched in any cluster (Figure 2, pct.min >0.01, log_2_ fold-difference >0.25) were identified using the ‘FindAllMarkers’ function in Signac. TF motif scores (Figure 3) were calculated using the R package chromVAR implemented in Signac (Ref. ^97^) and Homer v4.11 (Ref. ^98^). Enhancers were defined as regions >-1 kb and >2 kb from transcription start sites. ATAC-seq heatmaps (Figures 2, 3, 7, and S4, peak summits ±3 kb) were produced using the computeMatrix and plotHeatmap functions in DeepTools v3.3.2.0.170 (Ref. ^95^).

### Pseudotime trajectories, RNA velocity, and differential cell type abundance

Differentiation trajectories were computed from scRNA and snATAC data using Slingshot v2.4.0 (Ref. ^30^) after reducing dimensionality and clustering like cells. Pre-computed cell embedding and clusters from the Seurat pipeline served as the input function, with ISCs (assigned from specific gene expression – Figure S1D) as the starting cluster (root node). The “wrapper” function in Slingshot constructed a minimum spanning tree (MST) between nodes, then generated lineage inference curves. Using these principal curves, we mapped individual pseudotimes along each branch of the secretory lineage. RNA velocity analysis (Figure S1) was performed using CellDancer,^99^ which uses a deep learning model to infer per-cell rate parameters for transcriptional and splicing kinetics for each gene. Differential abundance of cell type composition from snATAC-seq was tested using Milo^100^ to compare WT (n=2) and *Apc^-/-^* (n=3) samples. Cellular neighborhoods (Figure S5) were constructed using the KNN graph to give average neighborhood size of 1,000 cells. Differential abundance was tested for each neighborhood and those with significant differences (*P_adj_*<0.05) are plotted.

## QUANTIFICATION AND STATISTICAL ANALYSIS

Statistical analyses were performed using GraphPad Prism 9 or R (https://www.r-project.org/). Figure legends specify biological or technical replicates for each experiment. Statistical results are presented as averages, with individual values shown. Error bars represent the SEM.

Graphical illustrations were created at BioRender.com.

---

**Figure S1.**
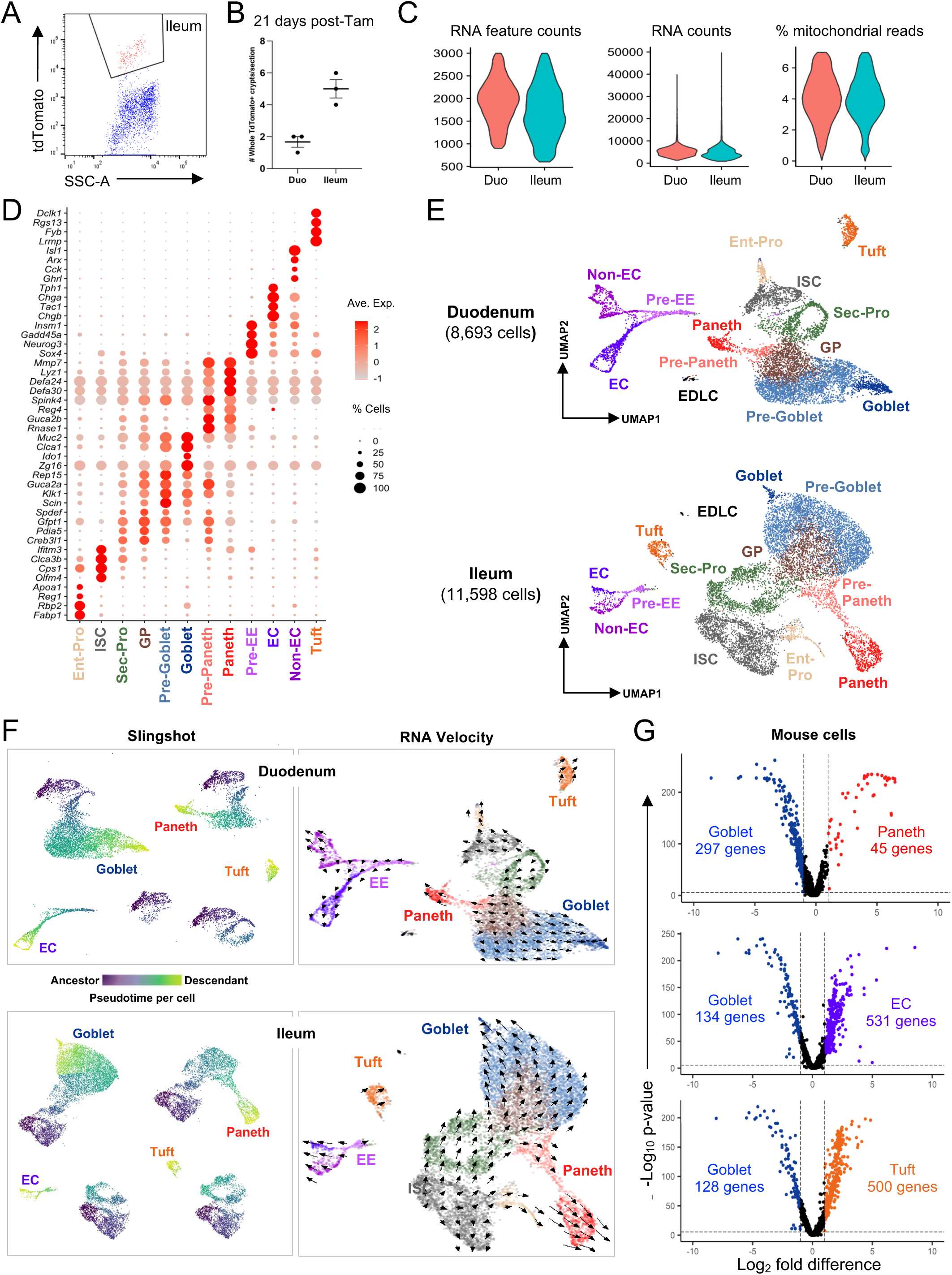
Distinct secretory cell populations labeled in *Atoh1^Cre-ER(T2)^;R26^Tom^* mice. Refers to Figure 1. **(A)** Representative FACs plot and gates to isolate Tom^+^ cells from ileal crypt epithelium. **(B)** Quantitation of clonal ribbons, which reflect lineage tracing from rare stem cells, 21 days after mice (n=3) received one dose of tamoxifen. **(C)** Quality control metrics from scRNA-seq of duodedal (Duo) and ileal Tom^+^ crypt cells. n=2 cell isolations from each region. **(D)** Expression of cell-specific markers enriched in each cluster, demonstrating capture of all expected cell types and accurate cluster assignments. **(E)** Separate UMAP plots from scRNA analysis of 8,693 duodenal and 11,598 ileal Tom^+^ cells, showing each cell population in both intestinal segments. n=2 mice for each segment. These data are combined in Figure 1C. **(F)** Pseudotime trajectories derived using Slingshot (left) or RNA velocity (right, cellDancer) for goblet, Paneth, EE, and tuft cells derived from ISCs and successive precursor states. Only cells that contribute to a trajectory acquire Slingshot scores (color) and dark-to-light gradients depict pseudotime progression. **(G)** Differentially expressed genes (FDR <0.05; log_2_ fold-difference >1, significance determined by the Wald test) between mouse intestinal goblet and other secretory cell types. Genes with the highest differential expression are listed.

**Figure S2.**
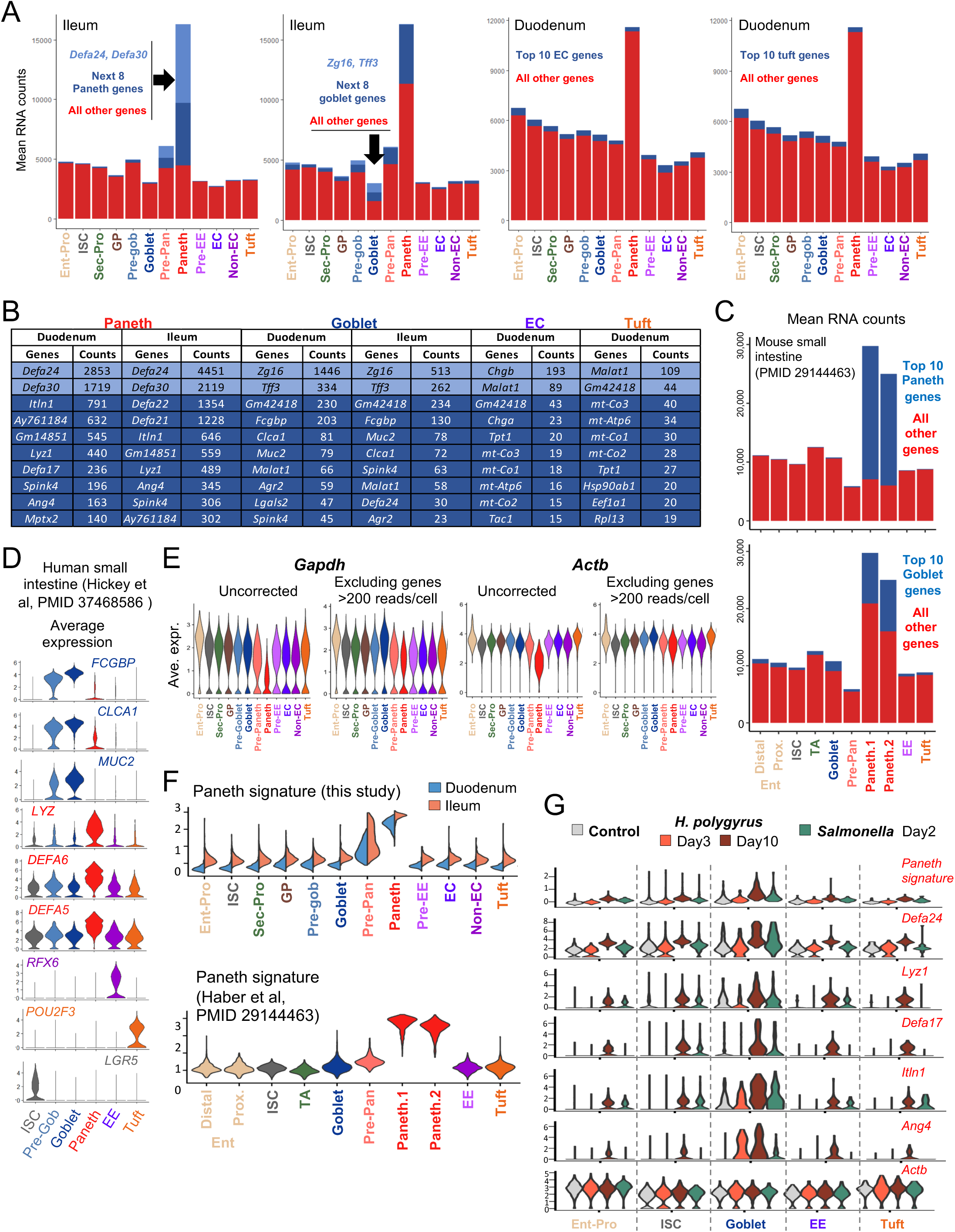
Paneth cell hyper transcription and shared TF genes with goblet cells. Refers to Figure 1. **(A)** Mean RNA counts from crypt cell populations, showing extreme values in Paneth cells and disproportionate enrichment of select genes in Paneth and goblet, compared to other secretory, cells. **(B)** Mean RNA counts of top 10 genes (by count) in Paneth, goblet, EC, and tuft cell clusters. The top 2 transcripts in Paneth and goblet cells have materially higher counts than the next 8. **(C)** Mean RNA counts in intestinal epithelial cell populations reported previously,^32^ affirming the extreme values we observed in Paneth cells. **(D)** Pseudobulk-averaged expression of conventional secretory-cell markers and *LGR5* in scRNA of human intestinal cells.^33^ Goblet and Paneth markers overlap in expression, while EE, tuft, and ISC markers are restricted. **(E)** Relative expression of ‘housekeeping’ genes *Gapdh* and *Actb* before and after corrections for extreme skewing from selected goblet and Paneth cell transcripts (removal of genes with mean RNA counts >200 from every cluster). **(F)** Signature of Paneth-enriched genes from Figure S1G shown in duodenal and ileal cell populations from this study (top) and signature of Paneth-enriched genes from scRNA-seq analysis in another study^32^ shown for each population in that study. **(G)** Pseudobulk-averaged expression of the Paneth enriched gene signature (from Figure S1G) and representative Paneth and housekeeping genes in wild type controls and helminth-(*H.polygyrus*) or *Salmonella*-infected mice.^32^ Paneth markers are enriched in goblet cells of infected animals.

**Figure S3.**
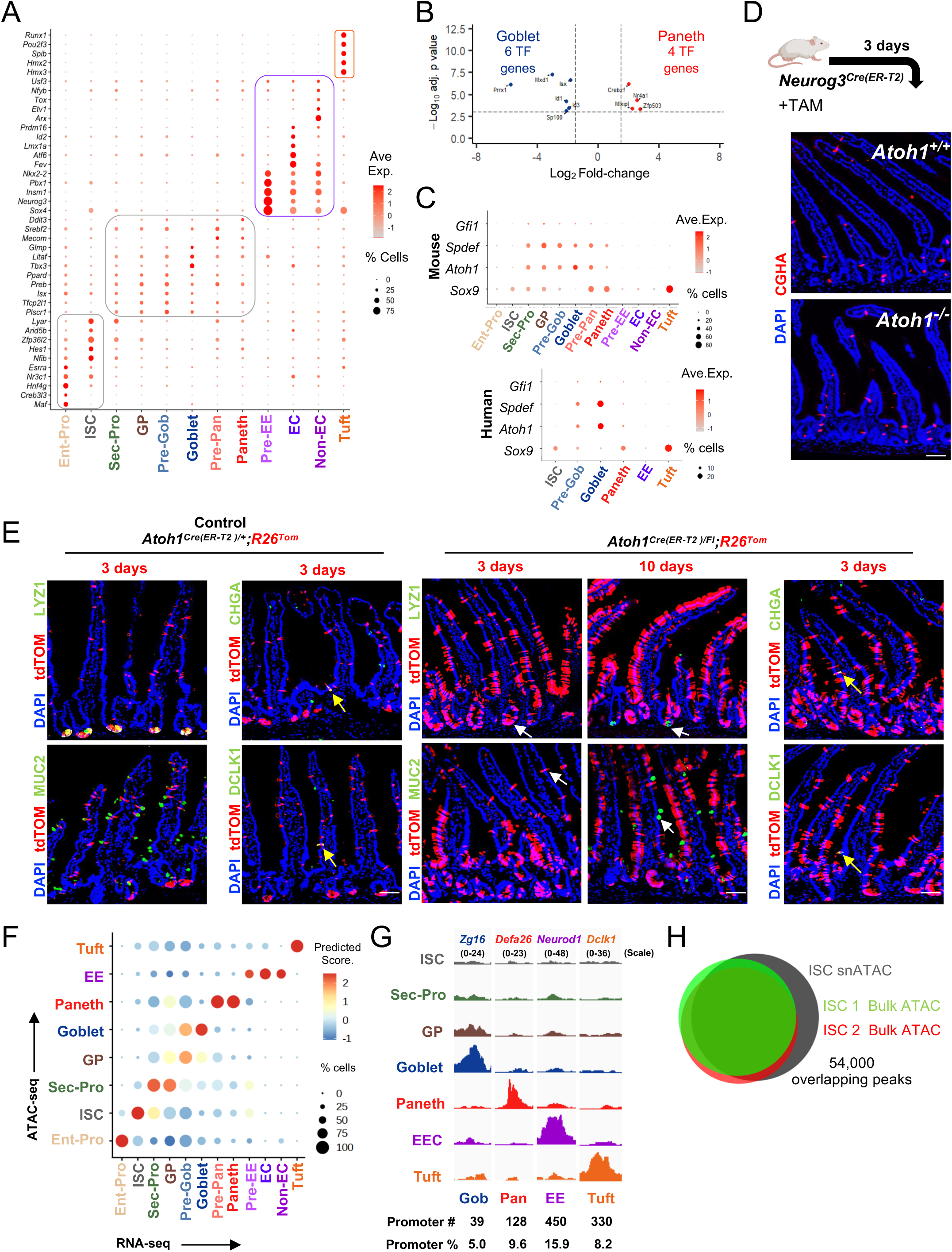
Limited TF diversity and *Atoh1* dependence of goblet and Paneth cells and epigenome of *Atoh1^Cre^*-labeled cells. Refers to Figure 2. **(A)** Top TFs enriched in each *Atoh1^Cre^* labeled cell cluster using the ‘FindAllMarkers’ function. Markers genes were selected based on minimum gene threshold of ‘pct.1’=0.25, meaning at least 25% of the cells in the cluster express those genes and a maximum threshold of ‘pct.2’= .30, meaning not more than 30% of cells in other clusters express the genes. In contrast to restricted TF expression in ISCs, enterocyte progenitors (Ent-Pro), EE, and tuft cells, goblet and Paneth cells show overlapping TFs and no unique enrichment. **(B)** TF genes differentially expressed (bulk RNA-seq, FDR <0.05; log_2_ fold-difference >1.5, significance determined by the Wald test) in mouse duodenal villus and crypt Tom^+^ cells isolated by flow cytometry 36 h or 12 days, respectively, after tamoxifen injection. **(C)** Relative expression of select TF genes reported as cell-specific or signal (Wnt)-responsive, in cell clusters from mouse (top) and human^33^ scRNA-seq. Circle diameter: within-cluster probability of gene detection, fill shade: normalized mean expression level. **(D)** Three days after tamoxifen treatment of *Neurog3^Cre-ER(T2)^;Atoh1^fl/fl^* mice, loss of *Atoh1* in EE cells did not affect CHGA^+^ cell numbers, indicating that *Atoh1* is not required to maintain established EE cells. n=3 independent experiments, scale bar 50 µm. **(E)** Control (*Atoh1^Cre/**+**^;R26^Tom^*) and *Atoh1^Sec-null^* (*Atoh1^Cre/**Fl**^;R26^Tom^*) intestines were harvested 3 or 10 days after tamoxifen injection. *Atoh1* loss resulted in the disappearance of characteristic mature MUC2^+^ goblet and LYZ1^+^ Paneth cells (white arrows), while CHGA^+^ EE and DCLK1^+^ tuft cells (yellow arrows) persisted. These findings underscore a role for ATOH1 in preserving goblet and Paneth cell properties after secretory identity is established. n=3 independent experiments, scale bar 50 µm. **(F)** Cell types among the KNN clusters derived from snATAC-seq were identified by matching uniquely accessible promoters with expression of the corresponding genes in scRNA-seq data (from Figure 1C). Although promoter accessibility generally varies less between cell types than enhancer accessibility, hence elevating noise when pairing snATAC and scRNA data, access at cell-specific promoters allowed unambiguous identification of cell types. **(G)** Representative IGV tracks (pseudobulk ATAC) at illustrative secretory gene promoters, showing restricted chromatin accessibility in cognate cells. Below: numbers of promoters with access restricted to each cell type and their representation among differentially enriched peaks. **(H)** The profile of open chromatin in ISCs from snATAC-seq of *Atoh1Cre^/+^;R26^Tom^* crypt cells overlaps significantly with that of open chromatin identified in bulk ATAC-seq analysis of purified Lgr5^+^ duodenal ISCs (n=2), further enhancing confidence in the data and in assigning the ISC designation to a prominent KNN cell cluster.

**Figure S4.**
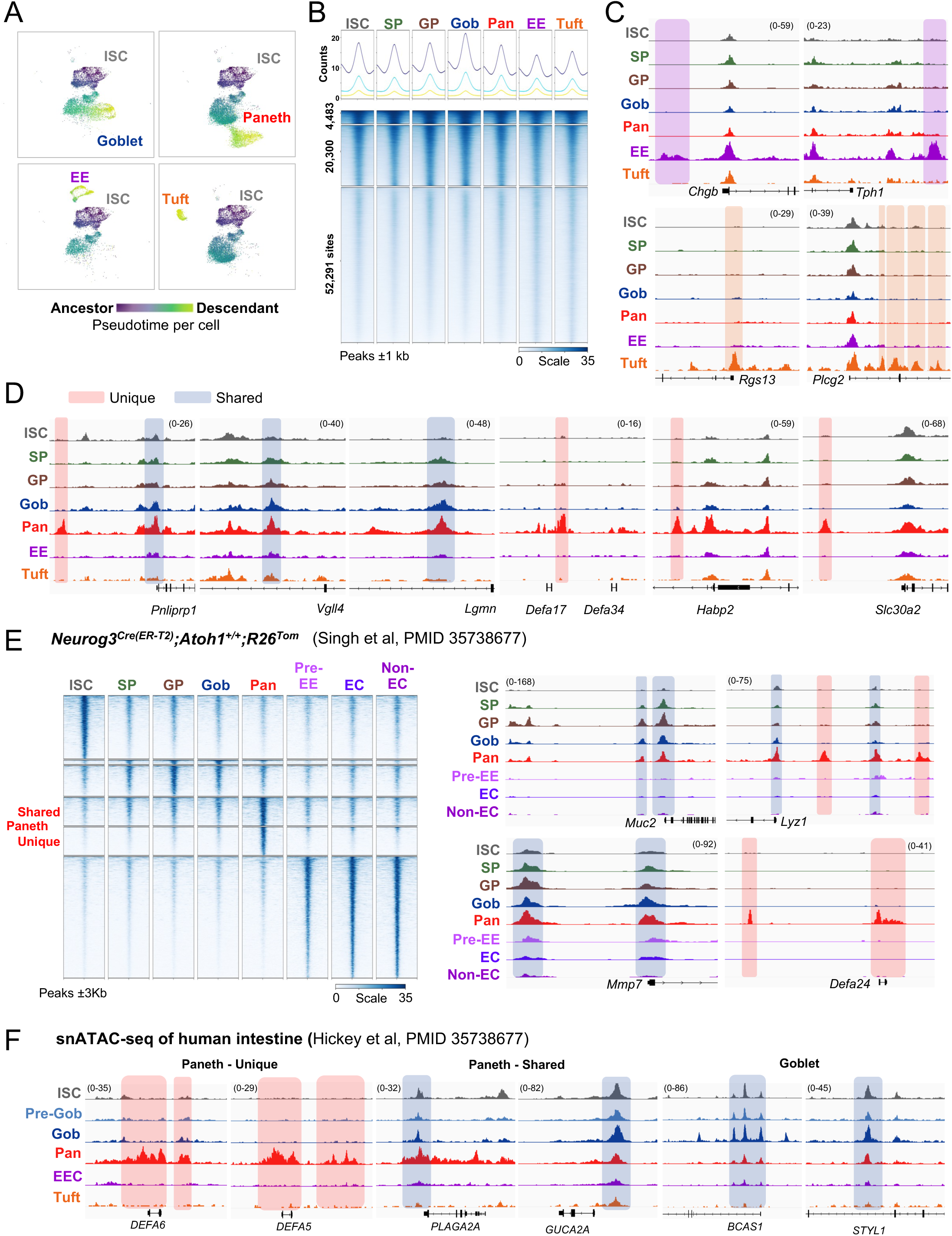
Shared and unique Paneth-cell *cis*-elements in *Atoh1^Cre-ER(T2)^;R26^Tom^* and *Neurog3^Cre-ER(T2)^;R26^Tom^* mice and in human intestinal cells. Refers to Figure 2. **(A)** Slingshot pseudotime trajectories derived from snATAC data, showing goblet, Paneth, EE, and tuft cells each originating in ISCs and successive secretory precursors. Only cells contributing to a *cis*-element trajectory acquire scores (color) and dark-to-light gradients depict pseudotemporal progression. **(B)** *k*-means clustering of pseudobulked ATAC-seq peaks enriched in >2 clusters. These signals show limited variance between cell populations. **(C-D)** Representative IGV tracks (pseudobulk ATAC) illustrating EE and tuft (**C**) and Paneth shared (accessible in goblet cells) and unique (inaccessible in other cells) *cis*-elements (**D**). Sites of note are shaded in colors corresponding to each secretory cell type. **(E)** Chromatin sites differentially accessible in distinct KNN cell clusters from *Atoh1^Cre-ER(T2)^;R26^Tom^* intestines, including shared and unique Paneth-cell *cis*-elements, were also differentially accessible in Tom^+^ cells isolated from *Neurog3^Cre-ER(T2)^;R26^Tom^* mice. Representative IGV tracks are shown below the heatmap. **(F)** Representative IGV tracks (pseudobulk from published snATAC data)^33^ from human Paneth and other intestinal epithelial cells, showing shared and unique Paneth and goblet *cis*-elements.

**Figure S5.**
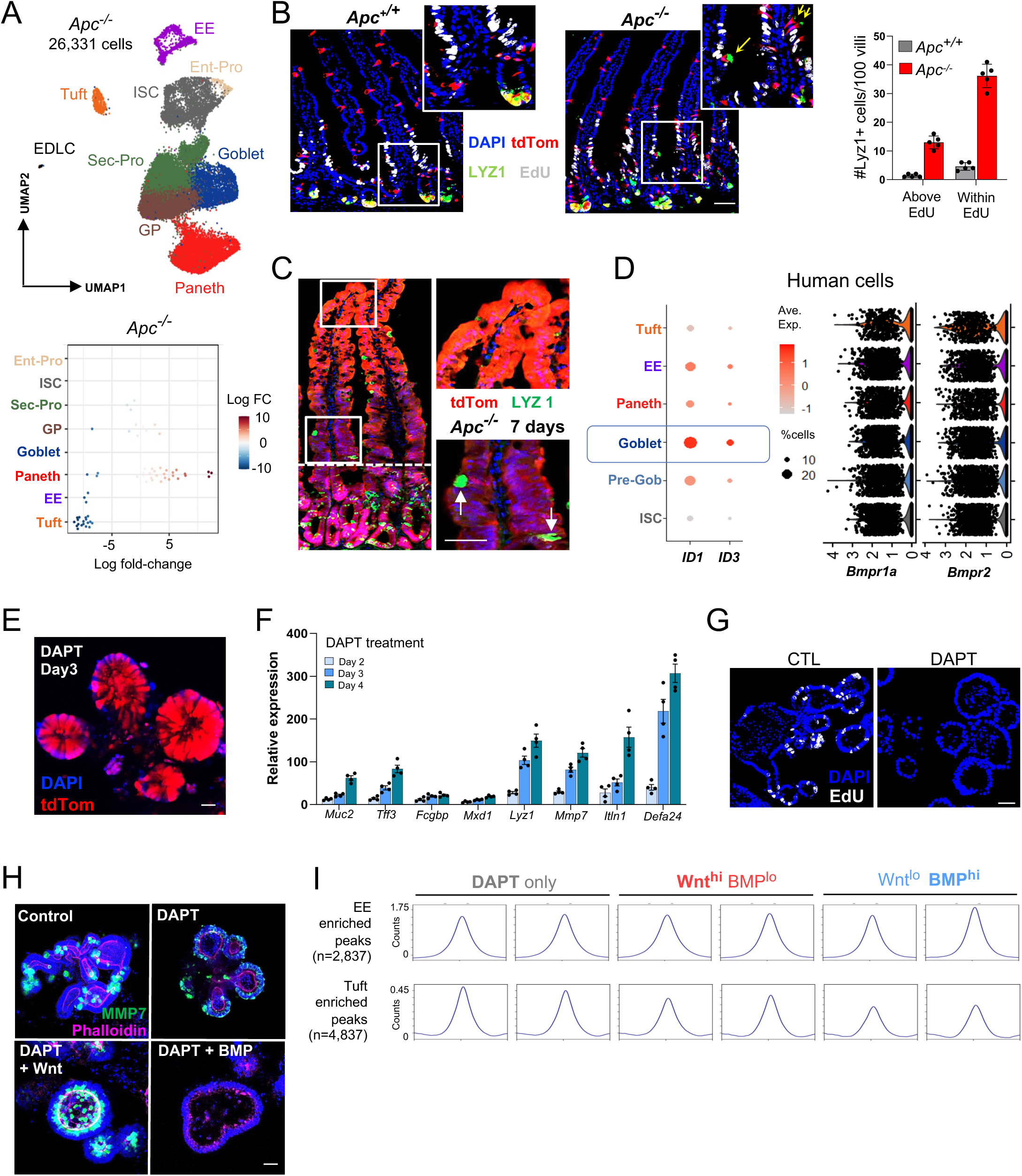
High Wnt/low BMP and high BMP/low Wnt response in mouse secretory cells. Refers to Figures 3, 4, and 5. **(A)** Integrated UMAP plots from snATAC analysis of duodenal Tom^+^ cells isolated from *Atoh1^Cre-ER(T2)^;R26^Tom^*;*Apc^fl/fl^* (26,331 cells merged from n=3 isolations). Cells were labeled with 2 doses of tamoxifen 24 h apart and harvested 12 h after the last dose. Cells with ATAC features of each population were recovered (top). Beeswarm plot depicting increased cells with Paneth features after forced Wnt activation (*Apc^-/-^*) in *Atoh1^Cre-ER(T2)^;R26^Tom^* mice. Cellular neighbourhoods were constructed with an average size of 1,000 cells and differential abundance was tested for each neighbourhood. Those with significant changes (FDR-adjusted p <0.05) are plotted. **(B)** Forced Wnt activation (*Apc^-/-^*) in *Atoh1^Cre-ER(T2)^;R26^Tom^* mice. Simultaneous single doses of tamoxifen and EdU distinguish pre-existing (above EdU column) from newly generated cells (within the EdU^+^ pool). LYZ1 and EdU immunostaining in intestines harvested 24 h later show increased numbers of ectopically positioned LYZ1^+^ cells, many with goblet morphology (inset), within and above the EdU column (quantified in the graph), reflecting a phenotypic shift from goblet to Paneth features. n=5 mice of each genotype, scale bars 50 µm. **(C)** Forced Wnt activation (*Apc^-/-^*) in *Villin^Cre-ER(T2)^;R26^Tom^* mice. A single dose of tamoxifen was administered to disrupt *APC* in intestinal epithelial cells and mice were harvested 7 days later. LYZ1 immunostaining showed increased numbers of ectopically positioned LYZ1^+^ cells, with goblet morphology largely confined to crypts, with LYZ1 detected in occasional mid-villus goblet cells (inset, white arrows) and absent on villus tip (inset). Scale bar 50 µm. **(D)** Human scRNA data^33^ showing enriched *ID1* and *ID3* expression in goblet cells (left, circle diameter: within-cluster probability of gene detection, fill shade: normalized mean expression level) and expression of BMP receptor genes in all epithelial cells (right: violin plots). **(E-F)** *Atoh1^Cre-ER(T2)^;R26^Tom^* mouse duodenal organoids were treated with DAPT to induce secretory differentiation. *Atoh1* induction was confirmed by presence of Tom^+^ organoids upon tamoxifen induction of Cre recombinase activity (**E**). RT-qPCR showed elevated expression of goblet and Paneth markers within 2 days of DAPT treatment and further increases thereafter (**F**). Scale bar 50 µm. **(G)** Mouse duodenal organoids were treated with or without DAPT for 3 days, grown in the presence of EdU for the last 30 min, and harvested to examine cell proliferation (EdU staining) which was absent in DAPT treated organoids. In the absence of new cell production, phenotypic changes after Wnt or BMP exposure would occur in existing cells. **(H)** Mouse duodenal organoids were treated with DAPT for 3 days, followed by Wnt^hi^ BMP^lo^ or Wnt^lo^ BMP^hi^ conditions for 4 days. In agreement with RT-qPCR findings, immunostaining revealed significantly increased MMP7 expression upon Wnt exposure and its absence upon BMP exposure. n=3 independent experiments, scale bar 50 µm. **(I)** Mouse duodenal organoids were treated with DAPT for 3 days, then cultured in Wnt^hi^ BMP^lo^ or Wnt^lo^ BMP^hi^ conditions for 4 additional days, and cells were collected for bulk ATAC-seq. In independent replicates, Rep, n=2 each), fragment counts reflecting accessible chromatin regions at EE- and tuft-enriched sites were largely unchanged in response to Wnt or BMP exposure.

**Figure S6.**
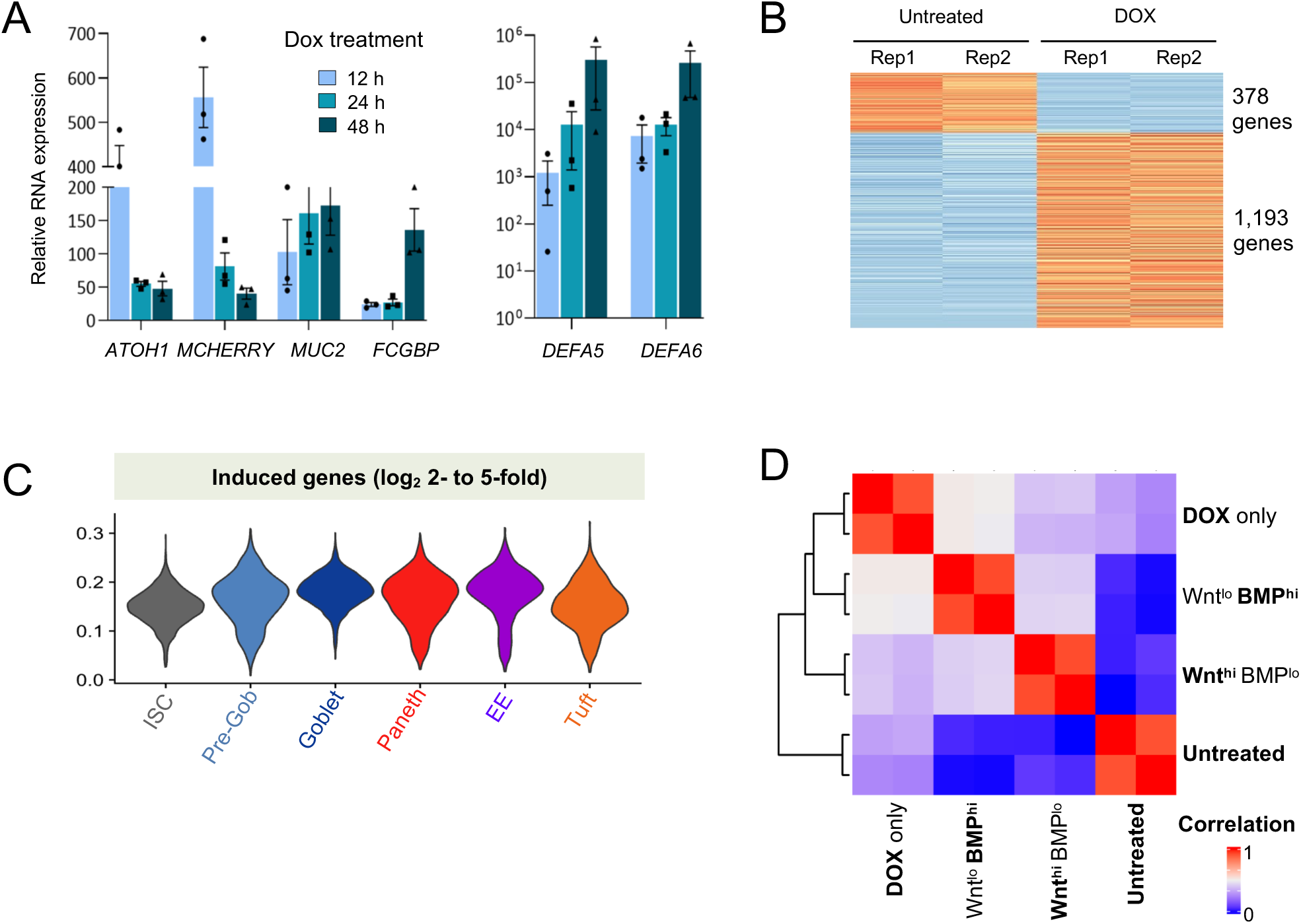
High Wnt/low BMP and high BMP/low Wnt response in human secretory cells. Refers to Figure 6. **(A)** RT-qPCR of *ATOH1* and conventional goblet and Paneth markers following Dox treatment of hISC^Atoh1(Dox)^ cells for 12, 24, and 48 h. Values are normalized to untreated (no Dox) cells. Goblet (left) and Paneth (right) genes are represented on different y-axis scales. n=3 experiments. **(B)** Genes differentially expressed in bulk RNA-seq analysis of independent replicates (Rep, n=2 each) of untreated and DOX-induced hISC^Atoh1(Dox)^ cells. **(C)** Distribution of transcripts enriched by log_2_ 2-to 5-fold upon Dox induction of hISC^Atoh1(Dox)^ cells (bulk RNA-seq, Figure 6A) in human intestinal epithelial cell populations (scRNAseq data from Ref. ^33^). Genes from each secretory cell type were induced in hISC^Atoh1(Dox)^ cells. **(D)** Correlation among duplicate bulk RNA-seq data from untreated hISC^Atoh1(Dox)^ cells and from cells treated only with Dox or, following Dox treatment, under Wnt^hi^ BMP^lo^ (CHIR 99021 + BMP inhibitor DMH1) or Wnt^lo^ BMP^hi^ (rBMP cocktail, no RSPO, no DMH1) conditions.

